# Harnessing FBXO31 with terminal amide-functionalized molecules for targeted protein degradation

**DOI:** 10.64898/2026.01.28.702440

**Authors:** Chenlu Zhang, Xiaokang Jin, Chen Zhou, M. Jamal Jenkins, Isabella A. Riha, Xiaoyu Zhang

**Affiliations:** Department of Chemistry, Northwestern University, Evanston, Illinois 60208, United States; Chemistry of Life Processes Institute, Northwestern University, Evanston, Illinois 60208, United States; Robert H. Lurie Comprehensive Cancer Center, Northwestern University, Chicago, Illinois 60611, United States; Center for Human Immunobiology, Northwestern University, Chicago, Illinois 60611, United States; International Institute for Nanotechnology, Northwestern University, Evanston, Illinois 60208, United States

**Author notes:** These authors contributed equally: Chenlu Zhang, Xiaokang Jin.

## Abstract

Targeted protein degradation (TPD) is a powerful strategy for controlling protein abundance. Here, we establish FBXO31 as a TPD-competent E3 ligase by exploiting its recognition of C-terminal amide-bearing degrons. Using an amidated Ala-Phe motif as a chemical recruiter, multiple small-molecule binders can be transformed into FBXO31-dependent degraders that induce rapid and potent target degradation. Mechanistic studies confirm FBXO31-mediated ternary complex formation and identify key residues in FBXO31 required for recruiter engagement and target degradation. We further show that an FBXO31-based multi-kinase degrader exhibits a distinct and broader degradation profile than a CRBN-based degrader, highlighting a potentially expanded degradable target space beyond CRBN.

## Main text

The ability to pharmacologically redirect cellular protein degradation pathways, including the proteasome^1-3^ and lysosome^4-6^, has emerged as a powerful strategy to modulate protein abundance for both basic research and therapeutic applications. This approach, known as targeted protein degradation (TPD), has thus far relied on a limited number of E3 ligases^7^, resulting in limited target coverage as well as context- and state-dependent degradation. Consequently, identifying and harnessing additional TPD-competent E3 ligases is essential for extending TPD to proteins with diverse subcellular localizations, expression levels, and biological contexts^8^. A recent study identified FBXO31, a substrate receptor of the SKP1-CUL1-F-box (SCF) ubiquitin ligase complex, as an E3 ligase that recognizes and promotes the proteasomal degradation of C-terminal amide-bearing proteins (CTAPs)^9^. This discovery adds to a growing number of N- and C-terminal degrons and their cognate E3 ligases^10^. In principle, these degron-E3 pairs offer attractive opportunities for heterobifunctional small-molecule degrader design by grafting terminal peptide degrons onto ligands that bind proteins of interest. However, many terminal degrons rely on charged or polar functional groups, which may compromise cell permeability and pose challenges for small-molecule degrader development. In this context, the FBXO31-C-terminal amide pair provides a distinctive and potentially advantageous platform, as recognition is enabled by a charge-neutral C-terminal amide. In addition, FBXO31 engagement prefers bulky and hydrophobic terminal residues, such as phenylalanine (Phe)^9^, which are suitable for incorporation into small-molecule degrader design with favorable drug-like properties.

Based on this rationale, we sought to explore the feasibility of harnessing a C-terminal amide as a chemical handle to functionalize small-molecule binders and convert them into degraders (**Figure 1a**). We first selected FKBP12 as a model substrate, as it has been used by our group and others due to its well-established degradability and the availability of a selective chemical probe, SLF (Synthetic Ligand of FKBP)^11-14^ (**Figure 1b**). Since an amidated C-terminal Phe was identified as the strongest binder to the FBXO31 degron-recognition pocket among all amino acids^9^, we initially synthesized four compounds by conjugating SLF to an amidated C-terminal Phe through different linkers (SLF-F1–F4, **Figure 1b**). To quantify induced FKBP12 degradation, we generated a C-terminal HiBiT-tagged^15^ FKBP12 at the endogenous locus in HEK293T cells, which also stably expressed HA-FBXO31 to enhance target degradation mediated by this ligase. As expected, FKBP12-HiBiT underwent dose-dependent degradation upon treatment with an established CRBN-based heterobifunctional degrader Len-SLF^16^ (**Figure 1c**). Using this system, we evaluated SLF-F1–F4 and found that three compounds, SLF-F2, SLF-F3, and SLF-F4, induced dose-dependent degradation of FKBP12-HiBiT, whereas SLF-F1 showed no detectable activity (**Figure 1c**). Among these compounds, SLF-F4 was the most potent, inducing FKBP12 degradation at concentrations as low as 5 nM and outperforming the positive control Len-SLF.

**Figure 1.**
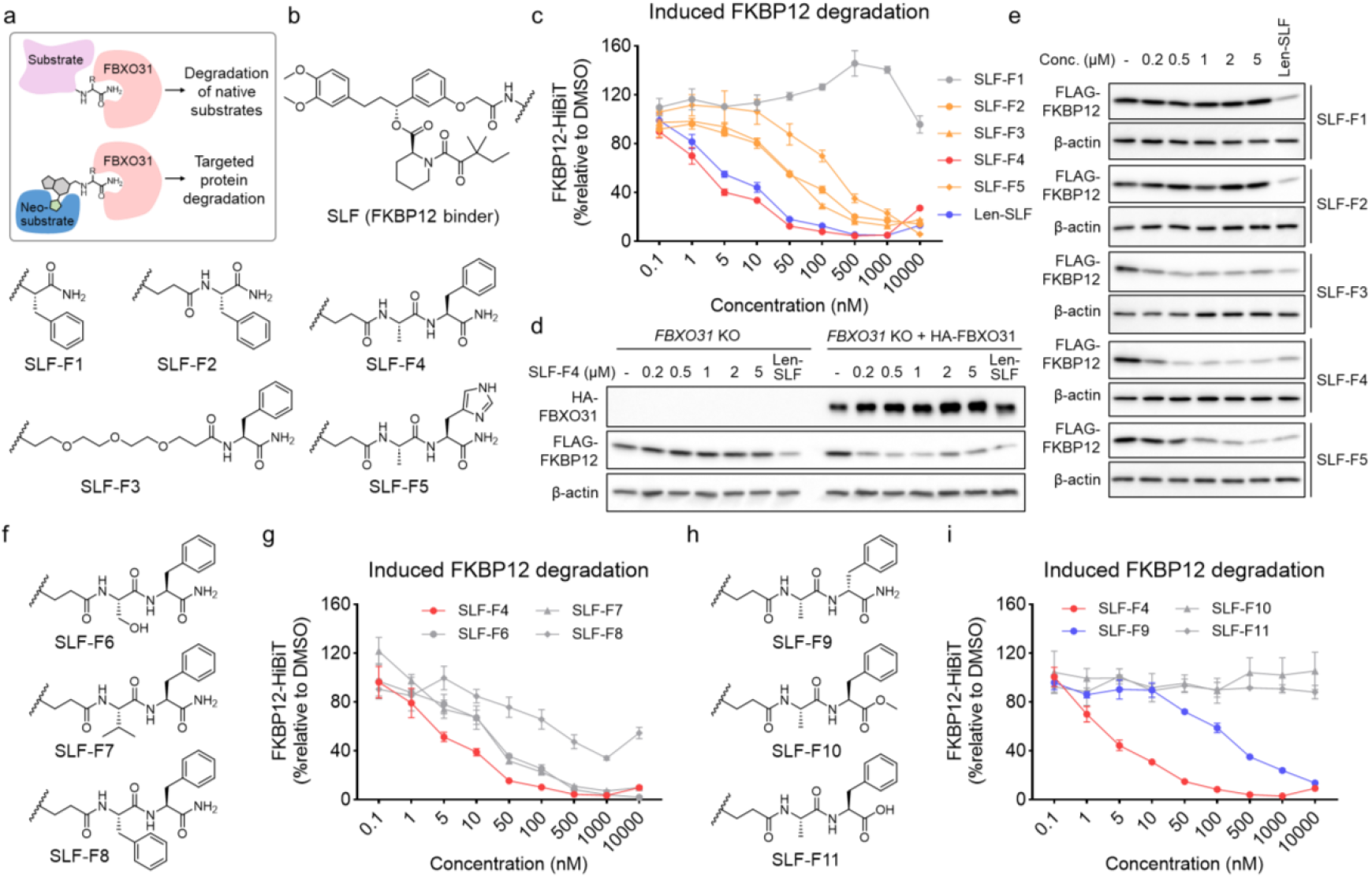
Design and validation of C-terminal amide-based heterobifunctional compounds engaging FBXO31 for FKBP12 degradation. **a**. Schematic illustrating the strategy of harnessing a C-terminal amide motif to recruit FBXO31 for TPD. **b**. Chemical structures of SLF-based FKBP12 degraders bearing amidated C-terminal Phe (SLF-F1– F4) or His (SLF-F5). **c**. Dose-response analysis of FKBP12-HiBiT degradation in HEK293T cells following treatment with SLF-F1–F5 or Len-SLF for 24 hours (n = 3). **d**. Western blot analysis of FLAG-FKBP12 levels in *FBXO31* KO HEK293T cells treated with SLF-F4 for 24 hours, with or without re-expression of HA-FBXO31. **e**. Western blot analysis of FLAG-FKBP12 levels in *FBXO31* KO HEK293T cells re-expressing HA-FBXO31 treated with SLF-F1–F5 for 24 hours. **f**. Chemical structures of SLF-F6–F8. **g**. Dose-response analysis of FKBP12-HiBiT degradation in HEK293T cells following treatment with SLF-F4 or -F6–F8 for 24 hours (n = 3). **h**. Chemical structures of SLF-F9– F11. **i**. Dose-response analysis of FKBP12-HiBiT degradation in HEK293T cells following treatment with SLF-F4 or -F9–F11 for 24 hours (n = 3).

To confirm that FKBP12 degradation was mediated by FBXO31, we generated *FBXO31* knockout (KO) HEK293T cells and further expressed FLAG-FKBP12 in these cells. Western blot analysis showed that SLF-F4, the most potent degrader, failed to degrade FLAG-FKBP12 in the absence of FBXO31 (**Figure 1d**), while re-introduction of HA- FBXO31 restored dose-dependent FKBP12 degradation by SLF-F4 (**Figure 1d**). Using this cell model, we further tested SLF-F1–F4 and observed degradation trends largely consistent with the HiBiT assay, with SLF-F4 remaining the most effective degrader and inducing FKBP12 degradation at the lowest tested concentration (0.2 µM) (**Figure 1e**). One exception was SLF-F2, which induced FKBP12 degradation in the HiBiT system but not by Western blot analysis. We speculate that the higher expression level of overexpressed FLAG-FKBP12 renders it less susceptible to degradation compared with endogenously expressed FKBP12-HiBiT.

Given that C-terminal histidine (His) provides the second strongest binding to FBXO31 degron-recognition pocket^9^, we replaced the amidated C-terminal Phe in the most potent degrader, SLF-F4, with an amidated C-terminal His to generate SLF-F5 (**Figure 1b**). SLF-F5 induced effective and dose-dependent FKBP12 degradation in both the HiBiT and Western blot assays, but with reduced potency compared to SLF-F4 (**Figure 1c**,**e**). SLF-F4 contains a dipeptide C-terminal motif with alanine (Ala) at the second-to-last position. To assess the effect of residue at this position on degradation activity, we synthesized three additional analogs containing serine (Ser) (SLF-F6), valine (Val) (SLF-F7), or Phe (SLF-F8) at this position (**Figure 1f**). All three compounds effectively promoted FKBP12 degradation, although none surpassed SLF-F4 in potency (**Figure 1g**). Next, we sought to synthesize three control compounds: one containing an amidated C-terminal D-Phe (SLF-F9), one with a C-terminal Phe methyl ester (SLF-F10), and one with a C-terminal Phe carboxylic acid (SLF-F11) (**Figure 1h**). Neither SLF-F10 nor SLF-F11 induced FKBP12 degradation at concentrations up to 10 µM (**Figure 1i**). Surprisingly, SLF-F9 exhibited no degradation activity below 50 nM but began to induce FKBP12 degradation at higher concentrations, ultimately achieving degradation efficiencies (>90%) comparable to SLF-F4 at 10 µM (**Figure 1i**). The ^1^H NMR spectra of SLF-F4 and SLF-F9 exhibit distinct chemical shift patterns in the high-field region, indicating that SLF-F9 is unlikely contaminated with SLF-F4. While further exploration of D-Phe in recruiter design is beyond the scope of this study, these results suggest a potential alternative strategy for FBXO31-based degrader design.

Next, we selected the most potent degrader, SLF-F4, to further characterize its ability to induce target degradation. We first determined the DC_50_ (nM) and D_max_ values of SLF-F4 at time points ranging from 2 to 30 hours (**Figure 2a**). SLF-F4 achieved a DC_50_ of 53.6 nM and 78% D_max_ at 2 hours, indicating rapid degradation of FKBP12 (**Figure 2a**,**b**). With extended treatment, both maximal degradation and potency increased, reaching a D_max_ of 97% at 24 hours with a DC_50_ of 7.4 nM (**Figure 2b**). Notably, for all time points tested, a similar hook effect was observed, with diminished degradation activity at concentrations above 1 µM in the FKBP12-HiBiT system (**Figure 2a**). SLF-F4-induced FKBP12 degradation was blocked by the proteasome inhibitor bortezomib (BTZ)^17^, the neddylation inhibitor MLN4924, which inhibits SCF ubiquitin ligase activity^18^, and SLF (**Figure 2c**). These results confirm that SLF-F4-mediated FKBP12 degradation proceeds through the proteasome and a Cullin-RING E3 ligase. To assess proteome-wide selectivity, we performed global proteomics in *FBXO31* KO HEK293T cells and in *FBXO31* KO cells re-expressing FBXO31, treated with either DMSO or SLF-F4. These analyses revealed selective and FBXO31-dependent degradation of FKBP12, with no obvious off-target degradation (**Figure 2d, Figure S1**, and **Table S1**). To further validate this observation in a different cellular context, we used HepG2 cells, which express relatively high levels of FBXO31^19^, and generated a *FBXO31* KO line. Global proteomics of parental and *FBXO31* KO HepG2 cells treated with SLF-F4 similarly demonstrated FBXO31-dependent degradation of endogenous FKBP12 (**Figure S2** and **Table S2**).

**Figure 2.**
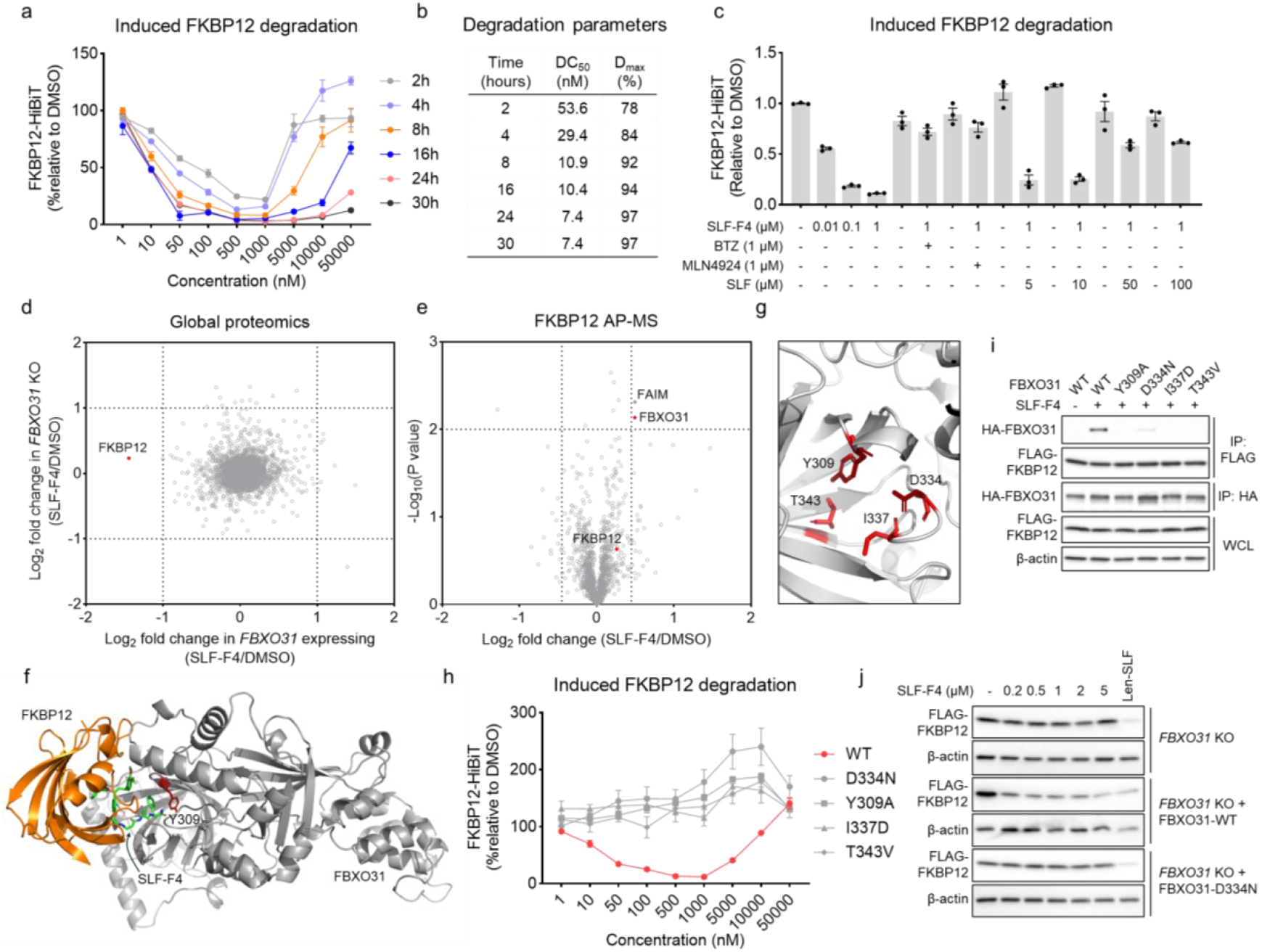
Mechanistic and proteomic characterization of SLF-F4-induced degradation of FKBP12. **a**. Dose-response curves for FKBP12 degradation induced by SLF-F4 at treatment times ranging from 2 to 30 hours in HEK293T cells (n = 3). **b**. DC_50_ and D_max_ values calculated from dose-response analyses in (a). **c**. Pharmacological rescue of SLF-F4-induced FKBP12 degradation by BTZ, MLN4924, and SLF (1 hour pretreatment followed by 7 hours of co-treatment.). **d**. Global proteomics analysis in *FBXO31* KO HEK293T cells and *FBXO31* KO cells re-expressing FBXO31 following SLF-F4 treatment (1 µM, 24 hours). **e**. AP-MS analysis of FLAG-FKBP12 from HEK293T cells treated with DMSO or SLF-F4 (1 µM, 4 hours). **f**. Ternary complex involving FKBP12, SLF-F4, and FBXO31 generated using the Boltz-2 model. The structure was rendered in PyMOL. **g**. Structural view of the FBXO31 degron-binding pocket (PDB: 5VZT) highlighting key residues (Y309, D334, I337, and T343) in substrate recognition. **h**. FKBP12-HiBiT degradation assays in *FBXO31* KO HEK293T cells re-expressing wild-type (WT) or mutant FBXO31 treated with SLF-F4 for 24 hours (n = 3). **i**. Co-immunoprecipitation in HEK293T cells expressing FLAG-FKBP12 and HA-FBXO31-WT or mutants, with or without SLF-F4 treatment (1 µM, 4 hours). All samples were treated with 0.1 µM BTZ. WCL, whole cell lysates. **j**. Western blot analysis of FLAG-FKBP12 levels in *FBXO31* KO HEK293T cells treated with SLF-F4 for 24 hours, with or without re-expression of HA-FBXO31-WT or -D334N.

To validate that SLF-F4 induces formation of a ternary complex involving FKBP12, the compound, and FBXO31, we performed affinity-purification mass spectrometry (AP-MS) using FLAG-FKBP12 enriched from HEK293T cells treated with either DMSO or SLF-F4. In the presence of SLF-F4, FKBP12 is associated with FBXO31, consistent with ternary complex formation (**Figure 2e** and **Table S3**). The SLF-F4-mediated neo-interaction is selective, as only one off-target interaction with FAIM was observed under the applied filtering criteria (Log2 fold change ≥ 0.45, -Log10 (P-value) ≥ 2). We next performed co-folding using the Boltz-2 model^20^, which yielded high-confidence property scores and revealed a well-formed ternary complex (**Figure 2f** and **Figure S3**). Since the interaction interfaces between SLF and FKBP12, as well as between the degron sequence and FBXO31, have been well characterized, this model enabled direct comparison with known structural features and showed strong agreement (**Figure S3**). For example, Y309 of FBXO31 forms a π-π stacking interaction with the C-terminal Phe side chain^9^, a feature accurately recapitulated in the co-folding model (**Figure 2f**). Previous studies identified four key residues within the FBXO31 degron-binding pocket, Y309, D334, I337, and T343, whose mutation disrupts substrate recognition and degradation^9^ (**Figure 2g**). We therefore expressed FBXO31-Y309A, -D334N, -I337D, or -T343V in *FBXO31* KO HEK293T cells and assessed FKBP12-HiBiT degradation by SLF-F4. Consistent with their loss-of-function properties, none of these FBXO31 variants supported FKBP12 degradation (**Figure 2h**). Importantly, these mutations are unlikely to cause major conformational defects, as all variants retained the ability to associate with SKP1, CUL1, and RBX1 in FBXO31-based AP-MS experiments (**Figure S4a** and **Table S4**). Thus, the loss of degradation activity is most consistent with disrupted interaction with the chemical recruiter, as validated by the co-immunoprecipitation experiment (**Figure 2i**). Next, using FBXO31-D334N as a representative mutant, we confirmed in an alternative FLAG-FKBP12-expressing system that SLF-F4 failed to induce FKBP12 degradation (**Figure 2j**). A similar loss of activity was also observed for another FKBP12 degrader SLF-F5 (**Figure S4b**).

To demonstrate the feasibility of harnessing FBXO31 to degrade additional protein substrates, we selected a multi-kinase binder TL13-87, which engages a large fraction of the kinome and has previously been used to generate heterobifunctional degraders for probing kinase degradation^21-23^. Based on the FKBP12 degradation study, we designed two degrader candidates incorporating the two most potent FBXO31 recruiters, resulting in MKI-F3 (**Figure S5a**) and MKI-F4 (**Figure 3a**). A previously reported CRBN-based degrader, SK-3-91, bearing the same kinase-binding scaffold, was included as a control compound (**Figure 3a**). To evaluate kinase degradation, we performed global proteomics in *FBXO31* KO HEK293T cells and *FBXO31* KO cells re-expressing FBXO31 (**Tables S5** and **S6**). While MKI-F3 showed no detectable kinase degradation activity (**Figure S5b**), MKI-F4, which incorporates the most potent recruiter, induced robust degradation of multiple kinases (**Figure 3b, Figure S6**, and **Table S7**). Consistent with FBXO31 dependence, degradation of all these kinases was abolished in *FBXO31* KO cells (**Figure 3b** and **Figure S6**). In contrast, the CRBN-based degrader SK-3-91 exhibited comparable kinase degradation profiles in both cell models (**Figure 3c** and **Figure S6**). Direct comparison of MKI-F4 and SK-3-91 revealed that MKI-F4 promotes degradation of a broader set of kinases, including several targets that were effectively degraded by MKI-F4 but were insensitive to SK-3-91 (**Figure 3d**). While more detailed comparisons are needed for future studies, these results highlight the potential of FBXO31 to enable degradation of substrates that are challenging to access using CRBN-based systems.

**Figure 3.**
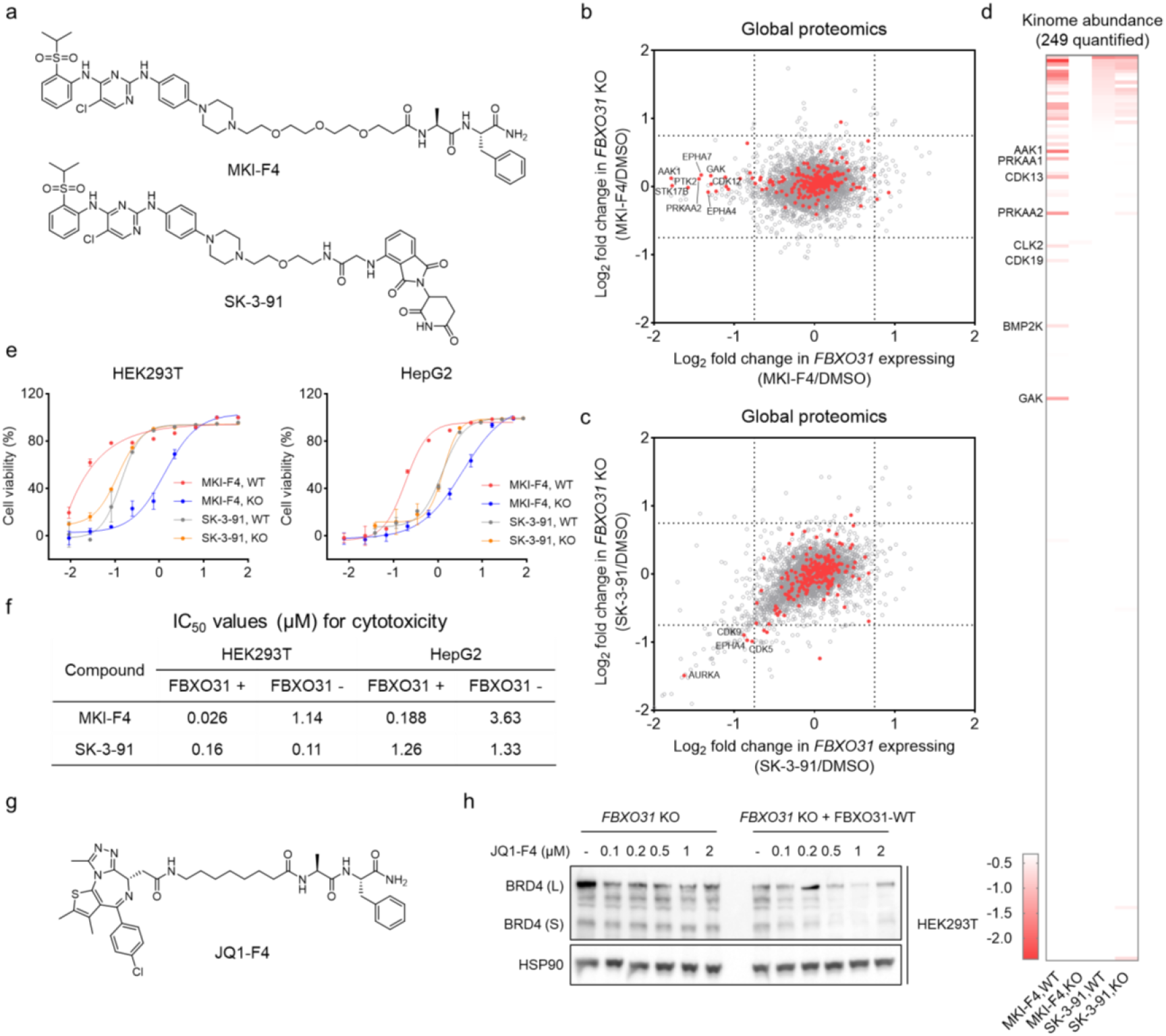
Harnessing FBXO31 to promote kinases and BRD4 degradation. **a**. Chemical structures of MKI-F4 and SK-3-91. **b**. Global proteomics analysis in *FBXO31* KO HEK293T cells and *FBXO31* KO cells re-expressing FBXO31 following MKI-F4 treatment (0.2 µM, 6 hours). Kinases are highlighted in red. **c**. Global proteomics analysis in *FBXO31* KO HEK293T cells and *FBXO31* KO cells re-expressing FBXO31 following SK-3-91 treatment (1 µM, 6 hours). Kinases are highlighted in red. **d**. Heatmap showing a direct comparison of kinase degradation profiles induced by MKI-F4 and SK-3-91 among the 249 quantified kinases. **e**. Dose-response curves showing MKI-F4-induced cytotoxicity in *FBXO31* KO HEK293T and HepG2 cells with or without FBXO31 re-expression (n = 3). **f**. Cytotoxic IC_50_ values of MKI-F4 and SK-3-91 in *FBXO31* KO cells and *FBXO31* KO cells re-expressing FBXO31. **g**. Chemical structure of JQ1-F4. **h**. Western blot analysis of BRD4 levels in *FBXO31* KO HEK293T cells and *FBXO31* KO cells re-expressing FBXO31 following JQ1-F4 treatment for 24 hours.

To assess the functional consequences of MKI-F4-mediated kinase degradation, we measured its cytotoxicity. Consistent with FBXO31-dependent activity, MKI-F4 displayed markedly reduced cytotoxicity in *FBXO31* KO cells, with IC_50_ values increasing 44-fold and 19-fold in HEK293T and HepG2 cells, respectively (**Figure 3e**,**f**). In contrast, both the inactive degrader MKI-F3 and the CRBN-based degrader SK-3-91 exhibited similar cytotoxicity profiles regardless of FBXO31 status (**Figure 3e,f** and **Figure S5c**). Finally, we synthesized a BRD4-targeting degrader, JQ1-F4 (**Figure 3g**), which induced dose-dependent BRD4 degradation in HEK293T and HepG2 cells expressing FBXO31, but not in *FBXO31* KO cells (**Figure 3h** and **Figure S7**). Collectively, these results establish FBXO31 and the amidated Ala-Phe motif as a potent recruitment strategy for degrading diverse protein substrates.

Introducing a minimal chemical handle provides a general and effective strategy for converting existing ligands into targeted protein degraders. To date, most such handles have been covalent or latent electrophiles that recruit E3 ligases, including DCAF16^24-27^, DCAF11^28^, and FBXO22^13, 29, 30^, through irreversible engagement of cysteine residues. In contrast, the FBXO31 recruitment strategy described here leverages a non-covalent chemical handle to drive target degradation, offering a complementary route for expanding the degrader toolbox.

## Acknowledgements

We gratefully acknowledge the support of the NIH R35GM154945 (X.Z.).

## Data Availability Statement

The mass spectrometry proteomics data have been deposited to the ProteomeXchange Consortium via the PRIDE^31^ partner repository with the dataset identifier PXD073606.

