## Supplementary Information for "Harnessing FBXO31 with terminal amide-functionalized molecules for targeted protein degradation"

### Supplementary Figures

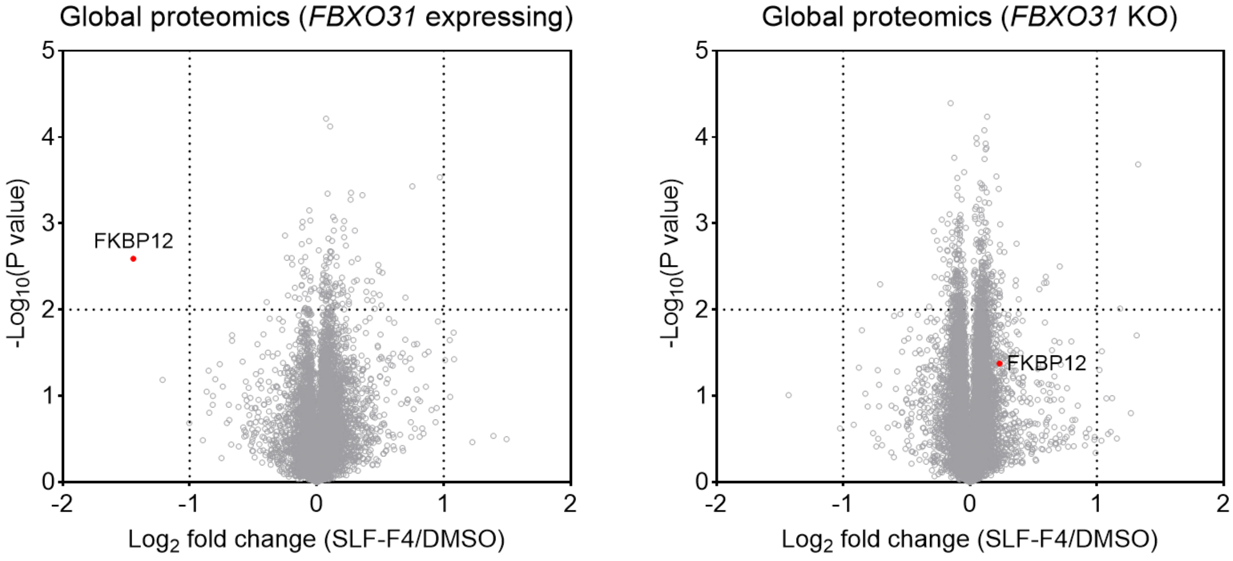


**Supplementary Figure 1. Proteomic profiling of SLF-F4-induced target degradation in HEK293T cells.** Volcano plots showing global proteome changes upon compound treatment in *FBXO31* KO HEK293T cells and *FBXO31* KO cells re-expressing FBXO31 (n = 3 biologically independent samples for SLF-F4; n = 2 biologically independent samples for DMSO). *P* values were calculated using a two-sided t-test and adjusted for multiple comparisons by the Benjamini-Hochberg method.

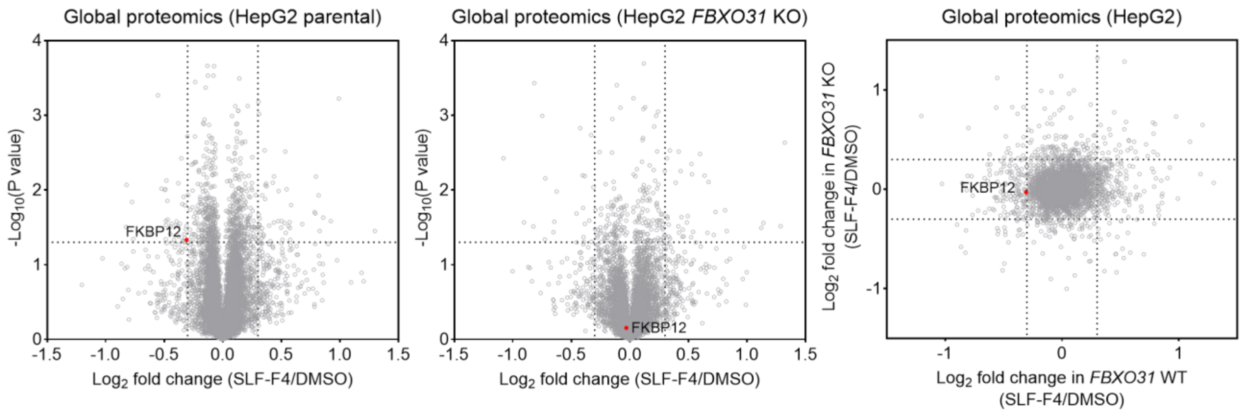


**Supplementary Figure 2. Proteomic profiling of SLF-F4-induced target degradation in HepG2 cells.** Volcano plots showing global proteome changes upon compound treatment in HepG2 parental and *FBXO31* KO cells (n = 3 biologically independent samples for SLF-F4; n = 2 biologically independent samples for DMSO). *P* values were calculated using a two-sided t-test and adjusted for multiple comparisons by the Benjamini-Hochberg method.

**Co-folding scores**

| Property | Value |
|---|---|
| Predicted TM-Score | 0.85 |
| ipTM | 0.895 |
| Confidence Score | 0.828 |
| Average pLDDT | 0.811 |
| Affinity Probability | 0.884 |
| Predicted pIC50 | 7.827 |
| Predicted IC50 | 14.9 nM |

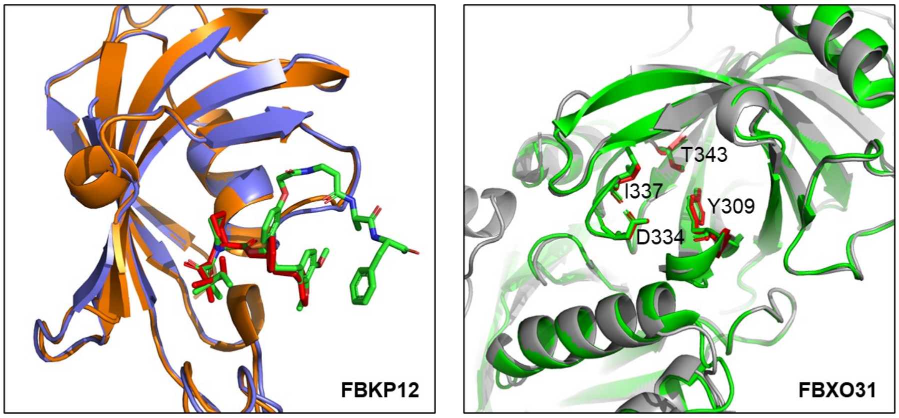


Orange: modeled FKBP12 bound to SLF-F4
Blue: FKBP12 bound to SB3 (red) (PDB: 1FKG)

Gray: modeled FBXO31 with key residues (red)
Green: FBXO31 with key residues (PDB: 5VZT)

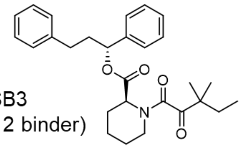

SB3
(FKBP12 binder)

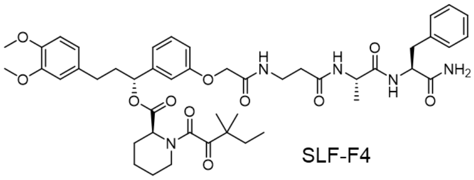

SLF-F4

**Supplementary Figure 3.** Co-folding scores and alignment of the modeled FKBP12-SLF-F4-FBXO31 ternary complex with reported crystal structures of FKBP12-ligand complex (PDB: 1FKG) and FBXO31 (PDB: 5VZT).

a

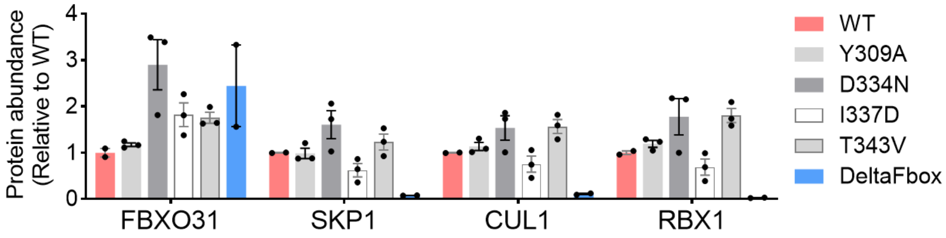


b

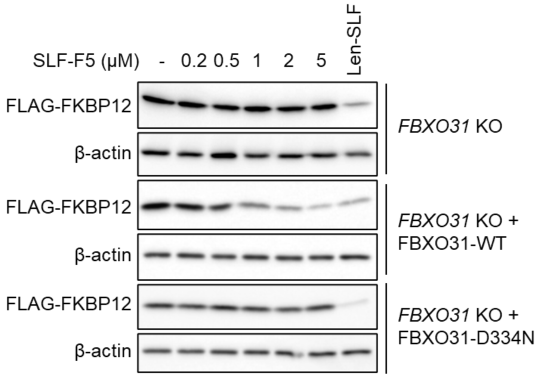


**Supplementary Figure 4. Functional analysis of key FBXO31 residues in target degradation. a**. AP-MS analysis shows that FBXO31-Y309A, -D334N, -I337D, and -T343V associate with SKP1, CUL1, and RBX1. FBXO31-DeltaFbox, which lacks the domain required for SKP1 interaction, fails to assemble into the complex. Data are presented as mean ± SEM (n = 3 biological independent samples for FBXO31-Y309A, -D334N, -I337D, and -T343V; n = 2 biological independent samples for FBXO31-WT and FBXO31-DeltaFbox). **b**. Western blot analysis of FLAG-FKBP12 levels in *FBXO31* KO HEK293T cells treated with SLF-F5 for 24 hours, with or without re-expression of HA-FBXO31-WT or -D334N.

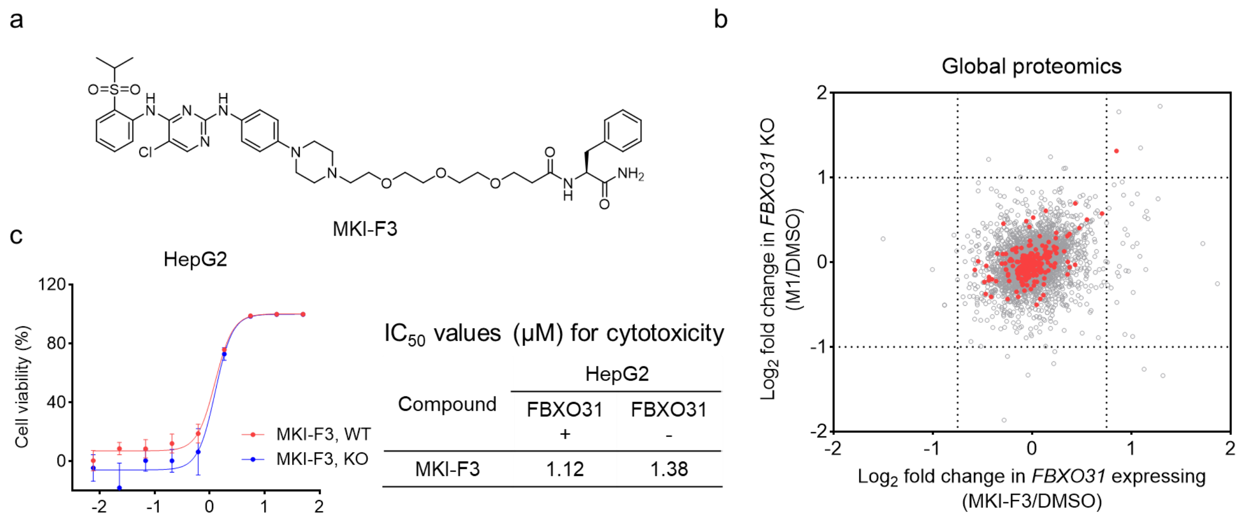


**Supplementary Figure 5. MKI-F3 shows no detectable kinase degradation. a.** Chemical structure of MKI-F3. **b**. Global proteomics analysis in *FBXO31* KO HEK293T cells and *FBXO31* KO cells re-expressing FBXO31 following MKI-F3 treatment (0.2 μM, 24 hours) (n = 3 biologically independent samples for MKI-F3; n = 2 biologically independent samples for DMSO). Kinases are highlighted in red. **c**. Dose-response curves and cytotoxic $IC_{50}$ values showing MKI-F3-induced cytotoxicity in *FBXO31* KO HepG2 cells with or without FBXO31 re-expression.

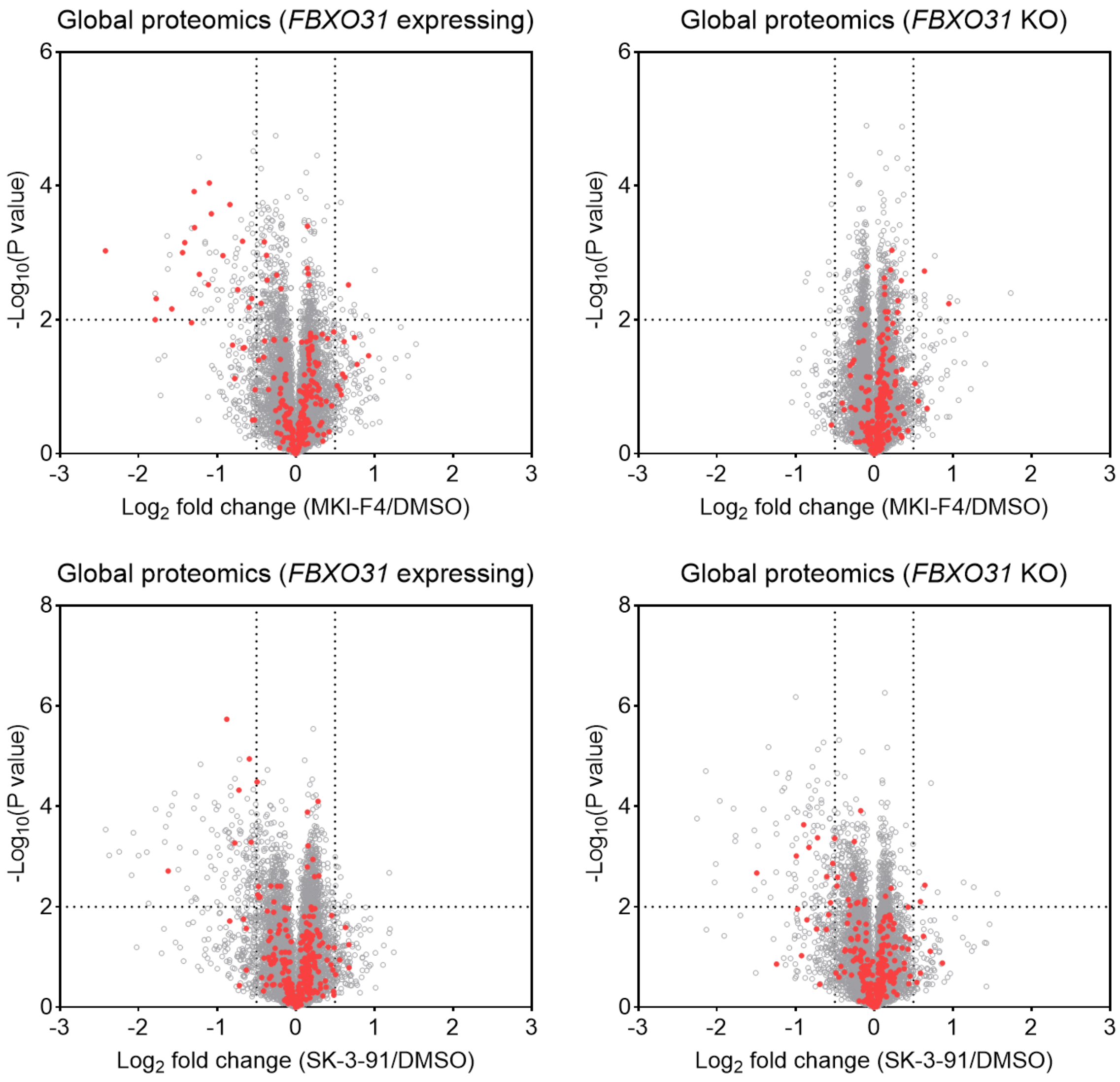


**Supplementary Figure 6. Proteomic profiling of MKI-F4- and SK-3-91-induced target degradation in HEK293T cells.** Volcano plots showing global proteome changes upon compound treatment in *FBXO31* KO HEK293T cells and *FBXO31* KO cells re-expressing FBXO31 (n = 3 biologically independent samples for MKI-F4 and SK-3-91; n = 2 biologically independent samples for DMSO). Kinases are highlighted in red. *P* values were calculated using a two-sided t-test and adjusted for multiple comparisons by the Benjamini-Hochberg method.

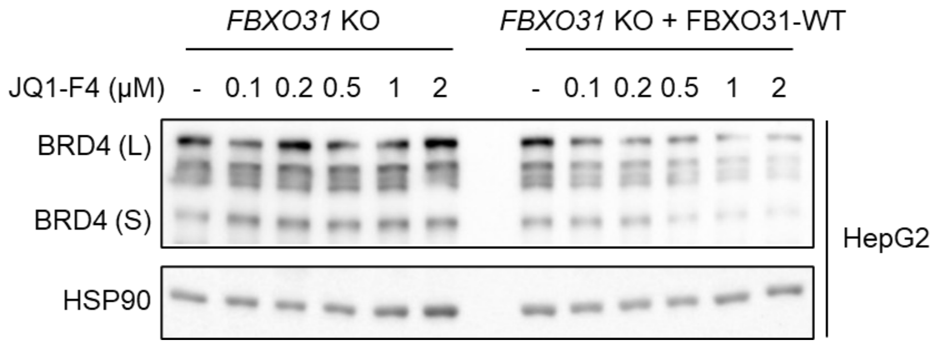


**Supplementary Figure 7.** Western blot analysis of BRD4 levels in *FBXO31* KO HepG2 cells and *FBXO31* KO cells re-expressing FBXO31 following JQ1-F4 treatment for 24 hours.

### Supplementary Materials and Methods

#### Reagents

The anti-HA (clone #C29F4, cat. #3724, dilution 1:5000), HRP-linked anti-HSP90 (clone #C45G5, cat. #79641, dilution 1:5000), HRP-linked rabbit IgG (cat. #7074, dilution 1:5000), and antibodies were purchased from Cell Signaling Technology. Anti-FBXO31 antibody (polyclonal, cat. #ab86137, dilution 1:1000) was purchased from Abcam. The anti-FLAG HRP antibody (clone M2, cat. #A8592, dilution 1:5000), anti-FLAG affinity gel (clone M2, cat. #A2220), anti-HA HRP antibody (clone #3F10, cat. #12013819001, dilution 1:5000), and cOmplete protease inhibitor cocktail (cat. #11873580001) were purchased from Sigma-Aldrich. The anti-β-Actin HRP antibody (clone #C4, cat. #sc-47778, dilution 1:1000) was purchased from Santa Cruz Biotechnology. Cas9 endonuclease was purchased from Integrated DNA Technologies (cat. #1081061). PureLink genomic DNA mini kit (cat. #K182001) was purchased from Invitrogen. FuGene 6 (cat. #E2692) transfection reagent, Nano-Glo HiBiT Lytic Detection System (cat. #N3040), Cell Titer Glo (cat. #G7573) and sequencing grade trypsin/Lys-C mix (cat. #V5071) were purchased from Promega. Blastcidin (cat. #ant-bl-05) was purchased from InvivoGen. Bortezomib (cat. #HY-10227) and SK-3-91 (cat. #HY-137341) were purchased from MedChemExpress. MLN4924 (cat. #15217) and SLF (cat. #10007974) were purchased from Cayman Chemical. DC protein assay kit (cat. #5000112) was purchased from Bio-Rad. Pierce high pH reversed-phase peptide fractionation kit (cat. #84868), Pierce ECL Western Blotting Substrate (cat. #32209), SuperSignal West Pico PLUS Chemiluminescent Substrate (cat. #34580) and Tandem Mass Tag (TMT) isobaric label reagent (cat. #90110) were purchased from Thermo Scientific.

#### Cell Lines

HEK293T cells were obtained from ATCC. HepG2 cells were obtained from The Developmental Therapeutics Core at Northwestern University. HEK293T and HepG2 cells were cultured in Dulbecco's Modified Eagle Medium (DMEM, Corning, cat.

#15013CV) with 10% (v/v) fetal bovine serum (FBS, Omega Scientific, cat. #FB-01) and L-glutamine (2 mM, Gibco, cat. #25030081). All the cell lines were tested negative for mycoplasma contamination using Universal Mycoplasma Detection Kit (ATCC, cat. #30-1012K).

**Cell Viability Assay**

HEK293T or HepG2 cells were seeded into a 96-well clear-bottom white plate (Corning, cat. #3610) at a density of 3,000 cells per well in 100 µL of DMEM. Subsequently, the cells were incubated with serial dilutions of test compounds, with a final DMSO concentration of 0.5% (v/v) in 100 µL of DMEM for 72 hours. Following treatment, 50 µL of Cell Titer Glo reagent (Promega, cat. #G7573) was added to each well and allowed to develop for 10 minutes at room temperature. Luminescence was measured using a CLARIOstar Plus microplate reader (BMG LABTECH).

**Generation of FKBP12-HiBiT Knock-in HEK293T cells**

Endogenous C-terminal HiBiT tagging of FKBP12 in HEK293T cells was achieved using CRISPR-Cas9-mediated homology-directed repair. The human FKBP12 genomic sequence was obtained from the Ensembl database and used to design a gRNA targeting a region proximal to the FKBP12 stop codon (FKBP12 gRNA: CATTTCCTTACCCAAGAACA). A single-stranded oligodeoxynucleotide (ssODN) donor template was designed to introduce the HiBiT tag upstream of the endogenous stop codon. The donor contained the HiBiT coding sequence followed by a stop codon, flanked symmetrically by 80-nucleotide homology arms corresponding to sequences upstream and downstream of the FKBP12 stop codon. The donor template sequence was as follows:

CCTATGGTGCCACTGGGCACCCAGGCATCATCCCACCACATGCCACTCTCGTCTTC
GATGTGGAGCTTCTAAAACTGGAAGGAGTGAGCGGCTGGCGGCTGTTCAAGAAGA
TTAGCTGACAGGAATGGCCTCCTCCCTTAGCTCCCTGTTCTTGGGTAAGGAAATGG
AATACTGAAGGGCCCTTCACTGCCTTTGCTCC

For genome editing, Cas9 ribonucleoprotein (RNP) complexes were assembled by incubating recombinant Cas9 protein with the gRNA at room temperature for 10 minutes. The assembled RNP complexes were then mixed with the ssODN donor template and delivered into HEK293T cells by electroporation. Successful FKBP12-HiBiT knock-in was confirmed using HiBiT detection assays upon treatment of DMSO or the FKBP12 degrader Len-SLF.

**HiBiT Assay**

FKBP12-HiBiT-expressing HEK293T cells were seeded into 96-well white plates at 5,000 cells/well and treated with tested compounds in triplicate. Following treatment, 50 µL of Nano-Glo HiBiT Lytic Detection System reagent (Promega, cat. #N3040) was added to each well and incubated for 10 minutes at room temperature. Luminescence was subsequently quantified using a CLARIOstar Plus microplate reader. In rescue experiments, cells were pretreated for 1 hour prior to the 7-hour co-incubation with tested compounds. For all experimental conditions, HiBiT luminescence signals were normalized to DMSO-treated control wells within each respective treatment group.

**Generation of *FBXO31* Knockout Cells**

*FBXO31* knockout HEK293T and HepG2 cells were generated by electroporating a complex consisting of Cas9 and three sgRNAs using the 4DNucleofector (Lonza Bioscience). Electroporation parameters were: pulse code CM-130 for HEK293T cells and EH-100 for HepG2 cells. Three sgRNAs targeting *FBXO31* gene were: sgRNA#1: UCUCATAGGGUCAUCGACGU; sgRNA#2: CCCGACGACCUCAUCAAGCC; sgRNA#3: GAGUGCAUGUACGGCCACAA. To confirm the knockout, genomic DNA was extracted using the PureLink Genomic DNA Mini Kit. *FBXO31* gene was amplified by Polymerase Chain Reaction (PCR) and measured via DNA gel electrophoresis and Sanger sequencing. The sequencing primers for *FBXO31* were: 5'-CTGGATTATTCTGCCTCCTCC-3' (forward) and 5'-ACTGCCAGACCAGATCAGAG-3' (reverse); 5'-GGTGACTTGGGCTCCTTTCT-3' (forward) and 5'-ACACAGACTCCCTGAGGGT-3' (reverse).

**Cloning, Mutagenesis, and Lentiviral Transduction**

The N terminal HA-tagged human FBXO31 in pCDH-CMV-MCS-EF1-Blast vector was obtained from GenScript. The FLAG-FKBP12 expression plasmid was constructed as previously described[1]. FBXO31 mutants (Y309A, D334N, I337D, T343V) were generated using the Gibson Assembly method (New England Biolabs). Lentiviral particles were produced by co-transfecting HEK293T cells with the target gene expression plasmid, psPAX2, and pMD2.G using FuGene 6 transfection reagent.
Viral supernatants were harvested 48 hours post-transfection, clarified through 0.45 μm Millex-HV sterile syringe filters (MilliporeSigma), and subsequently used to transduce HEK293T or HepG2 cells. Stable polyclonal cell lines were selected with blasticidin (10 μg/mL) for 7 days.

**Cell Lysis and Western Blot**

Cells were lysed in radioimmunoprecipitation assay (RIPA) buffer (25 mM Tris-HCl at pH 7.6, 150 mM NaCl, 1% NP-40, 1% sodium deoxycholate, and 0.1% SDS), supplemented with Universal Nuclease prior to use. The protein concentration was determined using the DC assay (Bio-Rad, cat. #5000112). The protein samples were mixed with Laemmli sample buffer, denatured by heating at 95 °C for 5 minutes. Proteins were separated using 4-20% Novex Tris-Glycine gels (Thermo Scientific, cat. #XP04205BOX) or 8% Novex Tris-Glycine gels (Thermo Scientific, cat. #XP00085BOX). Separated proteins were transferred to 0.2 μm PVDF membranes (Bio-Rad, cat. #1620177). Membranes were blocked with 5 % non-fat milk in TBST buffer (0.1% Tween-20, 20 mM Tris-HCl at pH 7.6, and 150 mM NaCl) for 1 hour at room temperature. Primary antibodies diluted in blocking buffer were applied with following conditions: FLAG, HA, β-Actin, and HSP90 antibodies were incubated for 1 hour at room temperature; other antibodies were incubated for 12 hours at 4 °C. Following three TBST washes, membranes were incubated with secondary antibody for 1 hour at room temperature. After three TBST washes, chemiluminescent signal was generated using ECL or Pico detection reagent and visualized on a ChemiDoc MP imaging system (Bio-Rad).

**Immunoprecipitations**

Cells were lysed in NP-40 lysis buffer (25 mM Tris-HCl at pH 7.6; 150 mM NaCl; 10% glycerol; 1% NP-40; cOmplete protease inhibitor cocktail). Cell lysate was centrifuged at 16,000 g for 10 minutes at 4 °C, and the resulting supernatant was used for subsequent immunoprecipitation. Protein lysates were added to FLAG affinity agarose (30 μL slurry per sample), which was pre-washed with immunoprecipitation washing buffer (25 mM Tris-HCl at pH 7.6; 150 mM NaCl; 0.2% NP-40), and rotated at 4 °C for 2 hours. The affinity agarose was then washed four times with immunoprecipitation washing buffer. The washed affinity agarose was mixed with Laemmli sample buffer and heated at 95 °C for 10 minutes. The resulting supernatant was used for western blot analysis.

**Global Proteomics**

Cells were lysed in 100 μL PBS containing cOmplete protease inhibitor cocktail using sonication (five pulses at 40% amplitude, repeated for three cycles). Total protein levels were quantified by DC protein assay. 100 μg of protein in 50 μL PBS was mixed with an equal volume of 12 M urea in PBS to achieve denaturation. Proteins were reduced by adding 5 μL of 200 mM DTT and incubating at 65 °C for 15 minutes, followed by alkylation with 5 μL of 400 mM iodoacetamide in water for 30 minutes at 37 °C in the dark. Samples were subsequently diluted with 300 μL PBS, and proteolytic digestion was performed using 2 μg trypsin/LysC at 37 °C for 12 hours. For TMT labeling, 8.5 μg of peptides were resuspended in 35 μL PBS, supplemented with 9 μL acetonitrile, and reacted with TMT reagents (5 μL per channel, 20 μg/μL in dry acetonitrile) for 1 hour at room temperature. Labeling reactions were terminated by sequential addition of 6 μL 5% hydroxylamine and 2.5 μL formic acid. Labeled samples were combined and fractionated into ten fractions using a high-pH reversed-phase peptide fractionation kit (Thermo Scientific, cat. #84868).

Eluted peptides were dried by vacuum centrifugation and analyzed by LC-MS on an Orbitrap Eclipse Tribrid mass spectrometer coupled with a Vanquish Neo UHPLC system.

Peptide samples were loaded onto an EASY-Spray analytical column (C18, 2 μm particles, 75 μm i.d., 150 mm length) and separated at a flow rate of 0.25 μL/min using a multistep gradient. Mobile phase A consisted of water containing 0.1% formic acid, and mobile phase B was 80% acetonitrile with 0.1% formic acid. The gradient was held at 5% B from 0-15 min, increased from 5% to 45% B over 15-155 min, and then ramped from 45% to 100% B between 155 and 180 min. Nanoelectrospray ionization was set at 1.5 kV. Data acquisition began with a full MS survey scan acquired in the Orbitrap (resolution 60,000; m/z 375-1600; RF lens 60%; standard AGC; automatic maximum injection time). For MS2, selected precursor ions were isolated in the quadrupole using a 0.7 m/z window and fragmented by higher-energy collisional dissociation in the ion trap (collision energy 27%; standard AGC; maximum injection time 35 ms). Following each MS2 event, up to ten fragment ions were selected by synchronous precursor selection (SPS) for MS3 analysis. These MS3 ions were further fragmented by HCD and detected in the Orbitrap (collision energy 55%; AGC target 250%; maximum injection time 200 ms; resolution 60,000). Raw spectra were acquired and recorded using Xcalibur software (v4.5.445.18, Thermo Scientific).

Raw mass spectrometry data were analyzed using Proteome Discoverer software (v2.5, Thermo Scientific). Peptide identification was performed with the Sequest HT algorithm by searching against the UniProt human reference proteome database (UP000005640_9606_Human.fasta). Searches used tryptic specificity, with mass tolerances set to 10 ppm for precursor ions and 0.6 Da for fragment ions. Carbamidomethylation of cysteines and TMT labeling on peptide N-termini and lysine residues were specified as fixed modifications. Methionine oxidation and protein N-terminal acetylation were specified as variable modifications. Peptide-spectrum matches were evaluated using Percolator, and identifications were filtered to a 1% false discovery rate at both the peptide and protein levels. Quantification of TMT reporter ions was extracted from SPS-MS3 scans using a 20 ppm integration window. Protein abundance values were calculated based on unique and razor peptides.

#### Affinity Purification Mass Spectrometry (AP-MS)

Cells were lysed in NP-40 lysis buffer containing 25 mM Tris-HCl at pH 7.6, 150 mM NaCl, 10% glycerol, 1% NP-40, and cOmplete protease inhibitor cocktail. Insoluble material was removed by centrifugation at 16,000 × g for 10 minutes at 4 °C, and the clarified lysates were used for immunoprecipitation. FLAG affinity agarose (30 μL slurry per sample) was added to the lysates and incubated with rotation for 2 hours at 4 °C. The agarose was washed four times with immunoprecipitation wash buffer, followed by two additional washes with PBS. Bound target and associated proteins were eluted by heating the agarose at 65 °C for 10 minutes in 8 M urea in PBS. Eluates were reduced with 12.5 mM DTT at 65 °C for 15 minutes and subsequently alkylated with 25 mM iodoacetamide at 37 °C for 30 minutes. Samples were diluted with PBS to lower the urea concentration to 2 M, followed by digestion with 2 μg of trypsin/LysC at 37 °C for 12 hours. TMT reagents (6 μL, in dry acetonitrile) were then added, and labeling was carried out at room temperature for 1 hour. Reactions were quenched by addition of 6 μL of 5% hydroxylamine and 2.5 μL of formic acid. Labeled peptides were combined, desalted using a Sep-Pak C18 cartridge (Waters, cat. #WAT054955), and dried by vacuum centrifugation. The resulting peptides were analyzed by LC-MS as described above. Data processing and quantification were performed using the same database search and analysis parameters described above.

#### Statistical Analysis

Quantitative data are presented as scatter plots, with error bars representing the mean ± standard error of the mean (SEM). For proteomics analyses, statistical significance was determined using two-sided Student's t-tests. Resulting *P* values were corrected for multiple comparisons using the Benjamini-Hochberg method.

### Synthesis of SLF-F1, SLF-F2 and SLF-F3

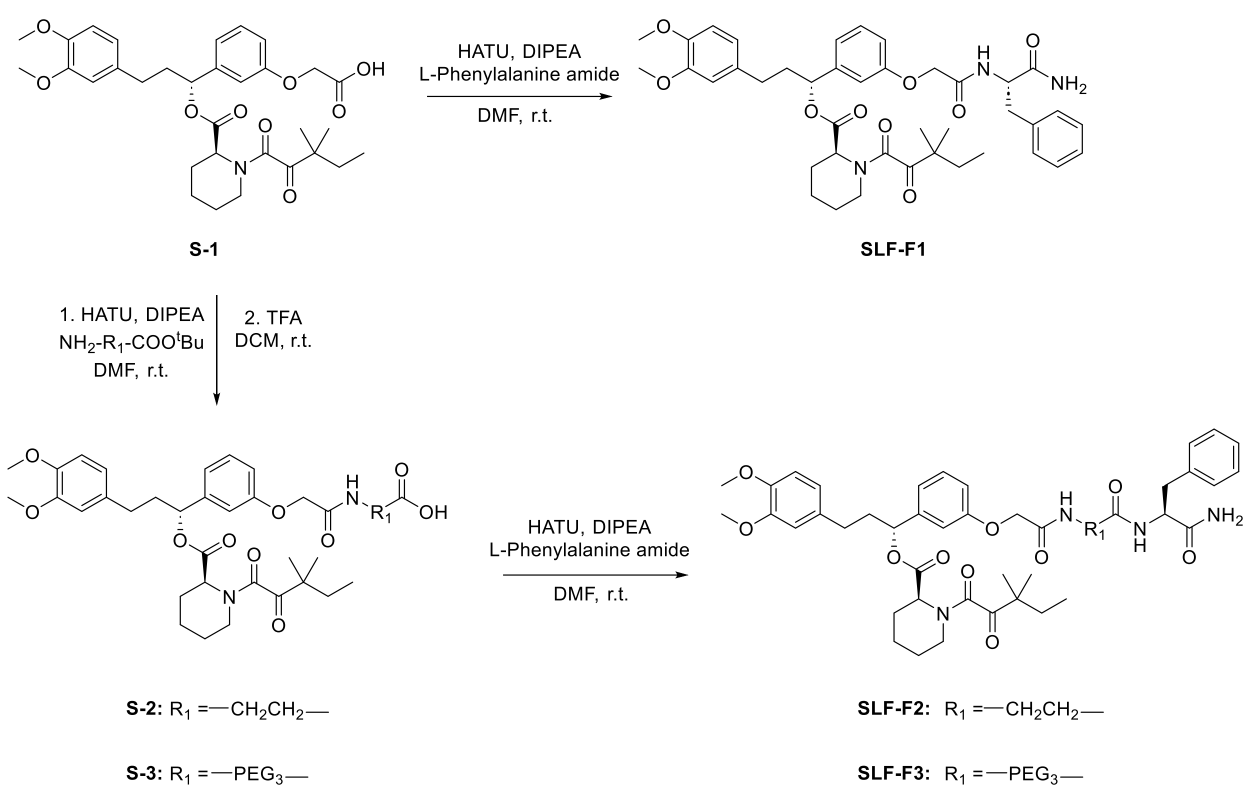


**SLF-F1:** *(R)-1-(3-(2-(((S)-1-amino-1-oxo-3-phenylpropan-2-yl)amino)-2-oxoethoxy)phenyl)-3-(3,4-dimethoxyphenyl)propyl (S)-1-(3,3-dimethyl-2-oxopentanoyl)piperidine-2-carboxylate*

S-1 was synthesized according to the previously reported method[2]. S-1 (50.0 mg, 0.086 mmol, 1.0 eq), HATU (42.6 mg, 0.112 mmol, 1.3 eq) and L-Phenylalanine amide (28.2 mg, 0.172 mmol, 2.0 eq) were dissolved in dry DMF (2.0 mL) at room temperature. DIPEA (71 µL, 0.430 mmol, 5.0 eq) was added to the reaction and stirred for 1 h. Upon completion, the reaction was concentrated, extracted with DCM (20 mL) and washed with brine (1 x 20 mL). The organic layer was dried by $Na_2SO_4$, concentrated, and the resulting residue purified by flash chromatography to provide SLF-F1 as white powder (45.2 mg, 0.062 mmol, 72%).

$^1$H NMR (500 MHz, Methanol-d$_4$) δ 7.30-7.27 (m, 1H), 7.23-7.14 (m, 5H), 7.02-6.98 (m, 1H), 6.94-6.92 (m, 1H), 6.86-6.82 (m, 2H), 6.80-6.77 (m, 1H), 6.72-6.70 (m, 1H), 5.77-5.72 (m, 1H), 5.23 (d, $J$ = 10 Hz, 1H), 4.73-4.70 (m, 1H), 4.54-4.43 (m, 2H), 3.80 (s, 3H), 3.79 (s, 3H), 3.42-3.35 (m, 1H), 3.18-3.13 (m, 2H), 3.01-2.97 (m, 1H), 2.66-2.54 (m, 2H), 2.35-2.33 (m, 1H), 2.29-2.20 (m, 1H), 2.10-2.02 (m, 1H), 1.77-1.59 (m, 5H), 1.52-1.27 (m, 2H), 1.21-1.10 (m, 6H), 0.89-0.77 (m, 3H). $^{13}$C NMR (126 MHz, Methanol-d$_4$) δ 209.03, 175.59, 171.00, 170.67, 169.14, 159.26, 150.47, 148.88, 143.39, 138.04, 135.24, 131.07, 130.42 (2C), 129.53 (2C), 127.91, 121.74, 121.13, 115.47, 114.37, 113.62, 113.29, 77.96, 68.19, 56.58, 56.47, 55.10, 52.85, 47.73, 45.74, 39.23, 39.07, 33.60, 32.27, 27.37, 25.91, 23.87, 23.64, 22.17, 9.13. HRMS (ESI$^+$) m/z calcd for $C_{41}H_{51}N_3NaO_9^+$ [M+Na]$^+$: 752.3523, found 752.3538.

**SLF-F2:** *(R)-1-(3-(2-((3-(((S)-1-amino-1-oxo-3-phenylpropan-2-yl)amino)-3-oxopropyl)amino)-2-oxoethoxy)phenyl)-3-(3,4-dimethoxyphenyl)propyl(S)-1-(3,3-dimethyl-2-oxopentanoyl)piperidine-2-carboxylate*

Step1: S-1 (60.0 mg, 0.103 mmol, 1.0 eq), HATU (50.9 mg, 0.134 mmol, 1.3 eq) and β-Alanine t-butyl ester hydrochloride (37.4 mg, 0.206 mmol, 2.0 eq) were dissolved in dry DMF (2.0 mL) at room temperature. DIPEA (85 μL, 0.515 mmol, 5.0 eq) was added to the reaction and stirred for 1 h. Upon completion, the reaction was concentrated, extracted with DCM (20 mL) and washed with brine (1 x 20 mL). The organic layer was dried by $Na_2SO_4$, concentrated, and the resulting residue was used without further purification. The crude product was dissolved in DCM (2.0 mL), TFA (0.5 mL) was added to the reaction and stirred for 2 h. The reaction mixture was then concentrated and purified by flash chromatography to provide S-2 as a colorless oil (56.2 mg, 0.086 mmol, 83.5% over 2 steps).

$^1$H NMR (500 MHz, $CDCl_3$) δ 7.70-7.65 (m, 1H), 7.31-7.28 (m, 1H), 6.98-6.93 (m, 2H), 6.84-6.78 (m, 2H), 6.71-6.67 (m, 2H), 5.84-5.72 (m, 1H), 5.41-5.33 (m, 1H), 4.67-4.55 (m, 2H), 3.86 (s, 3H), 3.85 (s, 3H), 3.72-3.62 (m, 2H), 3.41-3.38 (m, 1H), 3.29-3.23 (m, 1H),

2.65-2.54 (m, 4H), 2.46-2.43 (m, 1H), 2.27-2.21 (m, 1H), 2.12-2.05 (m, 1H), 1.85-1.62 (m, 5H), 1.57-1.38 (m, 2H), 1.25-1.18 (m, 6H), 0.91-0.83 (m, 3H).

Step2: S-2 (56.2 mg, 0.086 mmol, 1.0 eq), HATU (42.6 mg, 0.112 mmol, 1.3 eq) and L-Phenylalanine amide (28.2 mg, 0.172 mmol, 2.0 eq) were dissolved in dry DMF (2.0 mL) at room temperature. DIPEA (71 μL, 0.430 mmol, 5.0 eq) was added to the reaction and stirred for 1 h. Upon completion, the reaction was concentrated, extracted with DCM (20 mL) and washed with brine (1 x 20 mL). The organic layer was dried by $Na_2SO_4$, concentrated, and the resulting residue purified by flash chromatography to provide SLF-F2 as white powder (36.3 mg, 0.045 mmol, 52.3%).

$^1$H NMR (500 MHz, Methanol-d$_4$) δ 7.33-7.29 (m, 1H), 7.24-7.20 (m, 4H), 7.14-7.12 (m, 1H), 7.02-6.97 (m, 2H), 6.94-6.91 (m, 1H), 6.86-6.84 (m, 1H), 6.80-6.77 (m, 1H), 6.72-6.70 (m, 1H), 5.78-5.73 (m, 1H), 5.23 (d, *J* = 10 Hz, 1H), 4.65-4.62 (m, 1H), 4.46 (s, 2H), 3.80 (s, 3H), 3.79 (s, 3H), 3.49-3.38 (m, 3H), 3.22-3.13 (m, 2H), 2.88-2.83 (m, 1H), 2.66-2.54 (m, 2H), 2.42-2.23 (m, 4H), 2.10-2.03 (m, 1H), 1.77-1.61 (m, 5H), 1.53-1.33 (m, 2H), 1.22-1.11 (m, 6H), 0.89-0.78 (m, 3H). $^{13}$C NMR (126 MHz, Methanol-d$_4$) δ 209.05, 176.23, 173.53, 171.02, 170.93, 169.13, 159.26, 150.47, 148.89, 143.37, 138.64, 135.24, 131.02, 130.28 (2C), 129.43 (2C), 127.75, 121.74, 121.72, 121.07, 115.56, 114.50, 113.62, 113.30, 78.00, 68.22, 56.58, 56.48, 55.65, 52.85, 47.73, 45.74, 39.21, 39.02, 36.56, 36.27, 33.60, 32.27, 27.37, 25.93, 24.24, 23.87, 23.66, 22.17, 9.13. HRMS (ESI$^+$) m/z calcd for $C_{44}H_{56}N_4NaO_{10}^+$ [M+Na]$^+$: 823.3894, found 823.3897.

**SLF-F3:** *(R)-1-(3-(((S)-18-amino-17-benzyl-2,15,18-trioxo-6,9,12-trioxa-3,16-diazaoctadecyl)oxy)phenyl)-3-(3,4-dimethoxyphenyl)propyl (S)-1-(3,3-dimethyl-2-oxopentanoyl)piperidine-2-carboxylate*

The synthetic route of SLF-F3 was similar to SLF-F2.

$^1$H NMR (500 MHz, Methanol-d$_4$) δ 7.33-7.30 (m, 1H), 7.27-7.21 (m, 4H), 7.20-7.17 (m, 1H), 7.02-6.94 (m, 3H), 6.86-6.84 (m, 1H), 6.79-6.77 (m, 1H), 6.71-6.69 (m, 1H), 5.76-

5.73 (m, 1H), 5.22 (d, *J* = 10 Hz, 1H), 4.61-4.58 (m, 1H), 4.54 (s, 2H), 3.80 (s, 3H), 3.79 (s, 3H), 3.70-3.66 (m, 1H), 3.60-3.52 (m, 12H), 3.50-3.48 (m, 2H), 3.41-3.38 (m, 1H), 3.27-3.16 (m, 2H), 2.86-2.81 (m, 1H), 2.65-2.53 (m, 2H), 2.49-2.43 (m, 1H), 2.35-2.20 (m, 3H), 2.09-2.02 (m, 1H), 1.76-1.60 (m, 5H), 1.52-1.44 (m, 1H), 1.38-1.29 (m, 1H), 1.22-1.11 (m, 6H), 0.90-0.78 (m, 3H). $^{13}C$ NMR (126 MHz, Methanol-$d_4$) δ 209.05, 176.37, 174.16, 171.18, 171.00, 169.13, 159.41, 150.47, 148.88, 143.37, 138.93, 135.27, 131.02, 130.20 (2C), 129.48 (2C), 127.73, 121.75, 121.01, 115.78, 114.37, 113.64, 113.32, 78.03, 71.60, 71.53, 71.50, 71.44, 70.68, 68.69, 68.38, 56.59, 56.49, 55.86, 52.85, 47.73, 45.74, 40.07, 39.27, 38.35, 37.73, 33.61, 32.28, 27.38, 25.93, 23.87, 23.67, 22.17, 9.14. HRMS ($ESI^+$) m/z calcd for $C_{50}H_{68}N_4NaO_{13}^+$ $[M+Na]^+$: 955.4681, found 955.4686.

**Synthesis of SLF-F4, SLF-F5, SLF-F6, SLF-F7 and SLF-F8**

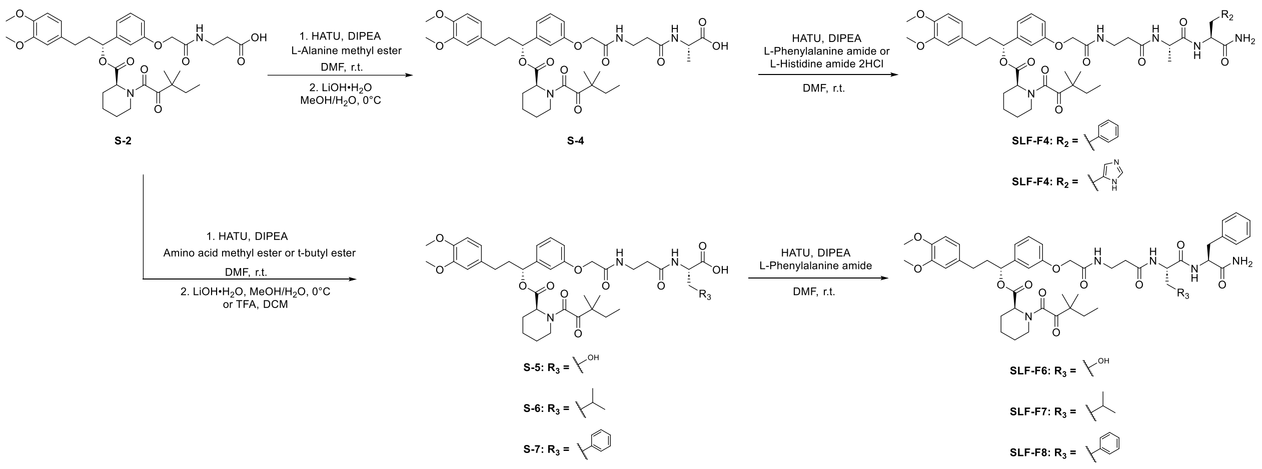


**SLF-F4:** *(R)-1-(3-(2-((3-(((S)-1-(((S)-1-amino-1-oxo-3-phenylpropan-2-yl)amino)-1-oxopropan-2-yl)amino)-3-oxopropyl)amino)-2-oxoethoxy)phenyl)-3-(3,4-dimethoxyphenyl)propyl (S)-1-(3,3-dimethyl-2-oxopentanoyl)piperidine-2-carboxylate*

Step1: S-2 (85.0 mg, 0.130 mmol, 1.0 eq), HATU (64.3 mg, 0.169 mmol, 1.3 eq) and L-Alanine methyl ester hydrochloride (36.3 mg, 0.260 mmol, 2.0 eq) were dissolved in dry DMF (3.0 mL) at room temperature. DIPEA (107 µL, 0.650 mmol, 5.0 eq) was added to the reaction and stirred for 1 h. Upon completion, the reaction was concentrated, extracted with DCM (30 mL) and washed with brine (1 x 30 mL). The organic layer was dried by $Na_2SO_4$, concentrated, and the resulting residue was used without further purification. The crude product was dissolved in MeOH/$H_2O$ (3.0 mL, 4:1), LiOH·$H_2O$(54.6 mg, 1.3 mmol, 10 eq) was added to the reaction and stirred at 0℃ for 0.5 h. 1M HCl was added to the reaction and pH was adjusted to 2, then concentrated and purified by flash chromatography to provide S-4 as a white powder (48.5 mg, 0.067 mmol, 51.5% over 2 steps).

$^1$H NMR (500 MHz, Methanol-$d_4$) δ 7.31 (t, $J$ = 8.0 Hz, 1H), 7.02-6.98 (m, 2H), 6.95-6.93 (m, 1H), 6.86-6.85 (m, 1H), 6.81-6.78 (m, 1H), 6.72-6.71 (m, 1H), 5.76-5.73 (m, 1H), 5.23-5.17 (m, 1H), 4.52 (s, 2H), 4.38-4.36 (m, 1H), 3.81 (s, 3H), 3.80 (s, 3H), 3.60-3.50 (m, 2H), 3.42-3.39 (m, 1H), 3.22-3.15 (m, 1H), 2.67-2.54 (m, 2H), 2.49-2.46 (m, 2H), 2.39-2.21 (m, 2H), 2.10-2.03 (m, 1H), 1.77-1.62 (m, 5H), 1.50-1.29 (m, 5H), 1.24-1.11 (m, 6H), 0.90-0.78 (m, 3H).

Step2: S-4 (23.0 mg, 0.032 mmol, 1.0 eq), HATU (15.7 mg, 0.041 mmol, 1.3 eq) and L-Phenylalanine amide (10.5 mg, 0.064 mmol, 2.0 eq) were dissolved in dry DMF (2.0 mL) at room temperature. DIPEA (27 μL, 0.160 mmol, 5.0 eq) was added to the reaction and stirred for 1 h. Upon completion, the reaction was concentrated, extracted with DCM (20 mL) and washed with brine (1 x 20 mL). The organic layer was dried by $Na_2SO_4$, concentrated, and the resulting residue purified by flash chromatography to provide SLF-F4 as white powder (14.8 mg, 0.017 mmol, 53.1%).

$^1$H NMR (500 MHz, Methanol-$d_4$) δ 7.23-7.16 (m, 1H), 7.16-7.09 (m, 5H), 6.92-6.87 (m, 2H), 6.85-6.82 (m, 1H), 6.77-6.74 (m, 1H), 6.70-6.67 (m, 1H), 6.62-6.60 (m, 1H), 5.67-5.63 (m, 1H), 5.12 (d, $J$ = 10 Hz, 1H), 4.49-4.45 (m, 1H), 4.42-4.40 (m, 2H), 4.10-4.05 (m, 1H), 3.70 (s, 3H), 3.69 (s, 3H), 3.48-3.38 (m, 2H), 3.31-3.28 (m, 1H), 3.12-3.06 (m, 2H), 2.90-2.85 (m, 1H), 2.56-2.44 (m, 2H), 2.39-2.31 (m, 2H), 2.25-2.23 (m, 1H), 2.19-2.11 (m, 1H), 1.99-1.92 (m, 1H), 1.66-1.51 (m, 5H), 1.41-1.19 (m, 3H), 1.13-1.01 (m, 8H), 0.79-0.68 (m, 3H). $^{13}$C NMR (126 MHz, Methanol-$d_4$) δ 209.05, 176.02, 174.90, 174.37, 171.10, 171.04, 169.14, 159.33, 150.47, 148.89, 143.38, 138.65, 135.26, 131.03, 130.39 (2C),

129.44 (2C), 127.75, 121.74, 121.04, 115.63, 114.45, 113.63, 113.30, 78.03, 68.31, 56.59, 56.48, 55.54, 52.86, 51.10, 47.73, 45.74, 39.23, 38.45, 36.70, 36.59, 33.61, 32.28, 27.37, 25.93, 23.86, 23.66, 22.17, 17.53, 9.13. HRMS (ESI$^+$) m/z calcd for $C_{47}H_{61}N_5NaO_{11}^+$ [M+Na]$^+$: 894.4265, found 894.4263.

The synthetic route of SLF-F5, SLF-F6, SLF-F7, SLF-F8 was similar to SLF-F4.

**SLF-F5:** *(R)-1-(3-(2-((3-(((S)-1-(((S)-1-amino-3-(1H-imidazol-4-yl)-1-oxopropan-2-yl)amino)-1-oxopropan-2-yl)amino)-3-oxopropyl)amino)-2-oxoethoxy)phenyl)-3-(3,4-dimethoxyphenyl)propyl (S)-1-(3,3-dimethyl-2-oxopentanoyl)piperidine-2-carboxylate*

$^1$H NMR (500 MHz, Methanol-d$_4$) δ 8.78-8.76 (m, 1H), 7.34-7.30 (m, 2H), 7.03-6.91 (m, 3H), 6.87-6.85 (m, 1H), 6.80-6.78 (m, 1H), 6.73-6.70 (m, 1H), 5.74-5.72 (m, 1H), 5.23-5.21 (m, 1H), 4.71-4.69 (m, 1H), 4.53 (s, 2H), 4.20-4.15 (m, 1H), 3.81 (s, 3H), 3.80 (s, 3H), 3.59-3.56 (m, 2H), 3.41-3.39 (m, 1H), 3.24-3.16 (m, 1H), 3.13-3.05 (m, 1H), 3.00-2.96 (m, 1H), 2.67-2.47 (m, 3H), 2.37-2.34 (m, 1H), 2.30-2.21 (m, 1H), 2.10-2.03 (m, 1H), 1.79-1.60 (m, 5H), 1.53-1.45 (m, 1H), 1.35-1.28 (m, 5H), 1.22-1.11 (m, 6H), 0.89-0.79 (m, 3H). $^{13}$C NMR (126 MHz, Methanol-d$_4$) δ 209.06, 175.37, 174.33, 174.10, 171.13, 171.05, 169.14, 159.26, 150.45, 148.87, 143.40, 135.20, 134.85, 131.06, 131.02, 121.72, 120.95, 118.53, 115.50, 114.46, 113.61, 113.28, 77.99, 68.31, 56.57, 56.47, 53.06, 52.86, 51.26, 47.70, 45.69, 39.22, 38.88, 36.67, 36.49, 33.57, 32.24, 27.80, 27.35, 25.89, 23.78, 23.66, 22.14, 9.09. HRMS (ESI$^+$) m/z calcd for $C_{44}H_{60}N_7O_{11}^+$ [M+H]$^+$: 862.4351, found 862.4345.

**SLF-F6:** *(R)-1-(3-(2-((3-(((S)-1-(((S)-1-amino-1-oxo-3-phenylpropan-2-yl)amino)-3-hydroxy-1-oxopropan-2-yl)amino)-3-oxopropyl)amino)-2-oxoethoxy)phenyl)-3-(3,4-dimethoxyphenyl)propyl (S)-1-(3,3-dimethyl-2-oxopentanoyl)piperidine-2-carboxylate*

$^1$H NMR (500 MHz, Methanol-d$_4$) δ 7.32-7.29 (m, 1H), 7.26-7.22 (m, 4H), 7.19-7.15 (m, 1H), 7.01-6.97 (m, 2H), 6.95-6.92 (m, 1H), 6.86-6.85 (m, 1H), 6.80-6.78 (m, 1H), 6.72-6.70 (m, 1H), 5.76-5.73 (m, 1H), 5.23-5.22 (m, 1H), 4.63-4.60 (m, 1H), 4.50 (s, 2H), 4.35-4.33 (m, 1H), 3.80 (s, 3H), 3.79 (s, 3H), 3.72-3.64 (m, 2H), 3.60-3.48 (m, 2H), 3.42-3.38

(m, 1H), 3.22-3.16 (m, 2H), 2.98-2.93 (m, 1H), 2.65-2.56 (m, 2H), 2.49-2.46 (m, 2H), 2.36-2.33 (m, 1H), 2.30-2.21 (m, 1H), 2.10-2.02 (m, 1H), 1.77-1.61 (m, 5H), 1.52-1.34 (m, 2H), 1.22-1.11 (m, 6H), 0.90-0.78 (m, 3H). $^{13}C$ NMR (126 MHz, Methanol-$d_4$) δ 209.05, 176.06, 174.28, 172.35, 171.10, 171.05, 169.15, 159.32, 150.48, 148.89, 143.37, 138.60, 135.26, 131.02, 130.36 (2C), 129.48 (2C), 127.78, 121.74, 121.04, 115.66, 114.45, 113.64, 113.31, 78.04, 68.30, 62.89, 56.99, 56.59, 56.48, 55.81, 52.87, 47.74, 45.75, 39.22, 38.46, 36.67, 36.51, 33.61, 32.28, 27.37, 25.93, 23.87, 23.66, 22.17, 9.13. HRMS (ESI$^+$) m/z calcd for $C_{47}H_{61}N_5NaO_{12}^+$ [M+Na]$^+$: 910.4214, found 910.4212.

**SLF-F7:** *(R)-1-(3-(2-((3-(((S)-1-(((S)-1-amino-1-oxo-3-phenylpropan-2-yl)amino)-3-methyl-1-oxobutan-2-yl)amino)-3-oxopropyl)amino)-2-oxoethoxy)phenyl)-3-(3,4-dimethoxyphenyl)propyl (S)-1-(3,3-dimethyl-2-oxopentanoyl)piperidine-2-carboxylate*

1H NMR (500 MHz, Methanol-$d_4$) δ 7.33-7.30 (m, 1H), 7.23-7.22 (m, 4H), 7.18-7.14 (m, 1H), 7.02-6.98 (m, 2H), 6.95-6.93 (m, 1H), 6.86-6.84 (m, 1H), 6.80-6.77 (m, 1H), 6.72-6.70 (m, 1H), 5.76-5.73 (m, 1H), 5.23-5.22 (m, 1H), 4.65-4.62 (m, 1H), 4.52 (s, 2H), 4.06-4.05 (m, 1H), 3.80 (s, 3H), 3.79 (s, 3H), 3.57-3.52 (m, 2H), 3.41-3.38 (m, 1H), 3.21-3.14 (m, 2H), 2.97-2.92 (m, 1H), 2.65-2.54 (m, 2H), 2.51-2.48 (m, 2H), 2.36-2.33 (m, 1H), 2.28-2.23 (m, 1H), 2.09-2.02 (m, 1H), 2.01-1.94 (m, 1H), 1.76-1.58 (m, 5H), 1.50-1.37 (m, 2H), 1.22-1.11 (m, 6H), 0.88 (t, $J$ = 7.5 Hz, 3H), 0.83-0.78 (m, 3H). $^{13}C$ NMR (126 MHz, Methanol-$d_4$) δ 209.04, 175.93, 174.45, 173.45, 171.04, 171.03, 169.14, 159.29, 150.48, 148.89, 143.39, 138.61, 135.25, 131.03, 130.38 (2C), 129.47 (2C), 127.74, 121.74, 121.07, 115.61, 114.48, 113.63, 113.31, 78.02, 68.32, 60.93, 56.59, 56.48, 55.57, 52.86, 47.73, 45.74, 39.23, 38.71, 36.79, 36.49, 33.61, 32.29, 31.46, 27.37, 25.93, 23.87, 23.67, 22.17, 19.61, 18.59, 9.13. HRMS (ESI$^+$) m/z calcd for $C_{49}H_{65}N_5NaO_{11}^+$ [M+Na]$^+$: 922.4578, found 922.4577.

**SLF-F8:** *(R)-1-(3-(2-((3-(((S)-1-(((S)-1-amino-1-oxo-3-phenylpropan-2-yl)amino)-1-oxo-3-phenylpropan-2-yl)amino)-3-oxopropyl)amino)-2-oxoethoxy)phenyl)-3-(3,4-dimethoxyphenyl)propyl (S)-1-(3,3-dimethyl-2-oxopentanoyl)piperidine-2-carboxylate*

$^{1}$H NMR (500 MHz, $CDCl_3$) δ 7.31-7.28 (m, 2H), 7.24-7.20 (m, 4H), 7.19-7.15 (m, 2H), 7.14-7.09 (m, 4H), 6.98-6.88 (m, 3H), 6.83-6.81 (m, 1H), 6.78-6.76 (m, 1H), 6.69-6.66 (m, 2H), 6.59-6.57 (m, 1H), 6.25-6.21 (m, 1H), 5.77-5.74 (m, 1H), 5.52-5.45 (m, 1H), 5.28 (s, 1H), 4.66-4.62 (m, 1H), 4.57-4.51 (m, 1H), 4.46-4.38 (m, 2H), 3.84 (s, 3H), 3.83 (s, 3H), 3.70-3.62 (m, 1H), 3.53-3.51 (m, 2H), 3.37-3.35 (m, 1H), 3.18-2.98 (m, 4H), 2.92-2.85 (m, 1H), 2.62-2.51 (m, 2H), 2.40-2.37 (m, 2H), 2.26-2.22 (m, 1H), 2.08-2.01 (m, 1H), 1.79-1.62 (m, 5H), 1.53-1.39 (m, 2H), 1.21-1.13 (m, 6H), 0.88-0.79 (m, 3H). $^{13}$C NMR (126 MHz, $CDCl_3$) δ 207.97, 173.11, 171.93, 170.86, 169.73, 168.80, 167.40, 157.40, 148.93, 147.41, 141.87, 136.64, 136.23, 133.36, 130.04, 129.34 (2C), 129.15 (2C), 128.78 (2C), 128.57 (2C), 127.16, 126.96, 120.20, 120.10, 114.07, 113.57, 111.78, 111.37, 76.60, 67.18, 55.95, 55.87, 51.38, 46.74, 44.20, 38.14, 37.72, 37.54, 35.78, 35.30, 32.48, 31.30, 26.47, 24.94, 23.45, 23.22, 21.20, 18.61, 17.27, 8.78. HRMS (ESI$^{+}$) m/z calcd for $C_{53}H_{65}N_5NaO_{11}^{+}$ [M+Na]$^{+}$: 970.4578, found 970.4577.

#### Synthesis of SLF-F9, SLF-F10 and SLF-F11

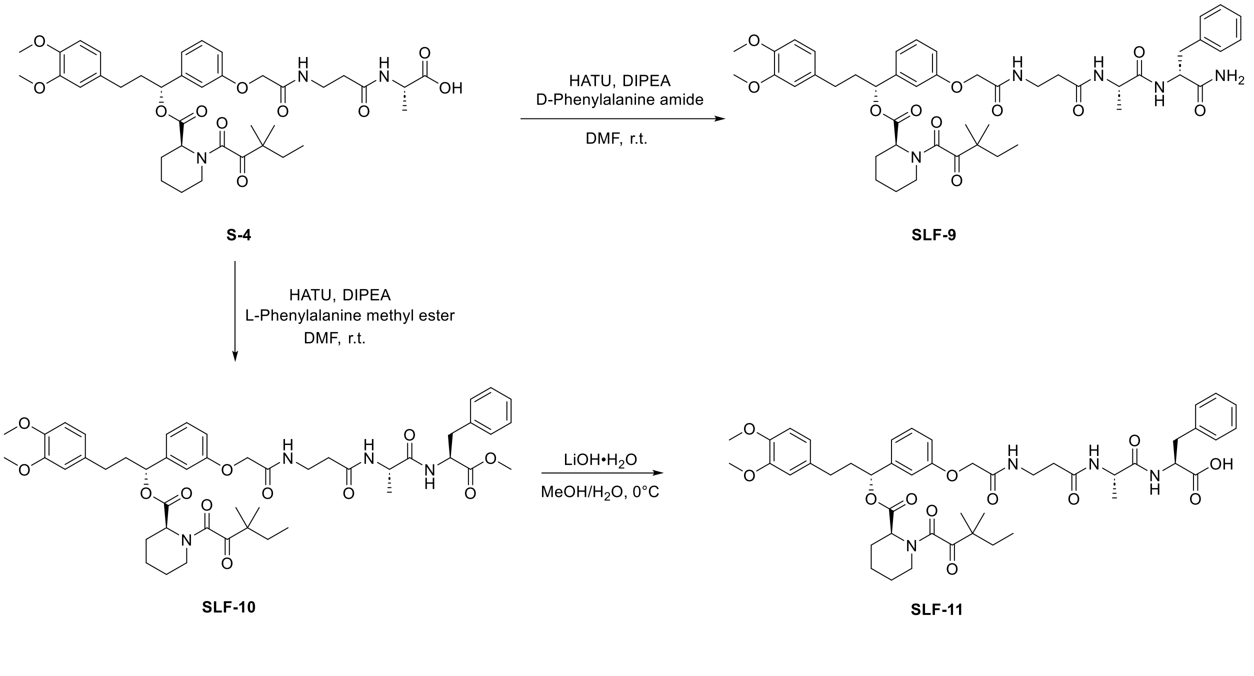


**SLF-F9:** *(R)-1-(3-(2-((3-(((S)-1-(((R)-1-amino-1-oxo-3-phenylpropan-2-yl)amino)-1-oxopropan-2-yl)amino)-3-oxopropyl)amino)-2-oxoethoxy)phenyl)-3-(3,4-dimethoxyphenyl)propyl (S)-1-(3,3-dimethyl-2-oxopentanoyl)piperidine-2-carboxylate*

S-4 (36.0 mg, 0.050 mmol, 1.0 eq), HATU (24.7 mg, 0.065 mmol, 1.3 eq) and D-Phenylalanine amide (16.4 mg, 0.10 mmol, 2.0 eq) were dissolved in dry DMF (2.0 mL) at room temperature. DIPEA (41 µL, 0.250 mmol, 5.0 eq) was added to the reaction and stirred for 1 h. Upon completion, the reaction was concentrated, extracted with DCM (20 mL) and washed with brine (1 x 20 mL). The organic layer was dried by $Na_2SO_4$, concentrated, and the resulting residue purified by flash chromatography to provide SLF-F9 as white powder (21.2 mg, 0.024 mmol, 48.0%).

$^1$H NMR (500 MHz, Methanol-d$_4$) δ 7.32-7.16 (m, 6H), 7.02-6.98 (m, 2H), 6.94-6.92 (m, 1H), 6.86-6.84 (m, 1H), 6.80-6.77 (m, 1H), 6.72-6.70 (m, 1H), 5.76-5.73 (m, 1H), 5.23-5.22 (m, 1H), 4.61-4.58 (m, 1H), 4.52-4.50 (m, 2H), 4.18-4.13 (m, 1H), 3.81-3.79 (m, 6H), 3.57-3.50 (m, 2H), 3.41-3.39 (m, 1H), 3.22-3.17 (m, 1H), 2.65-2.56 (m, 2H), 2.45-2.42 (m, 2H), 2.36-2.33 (m, 1H), 2.29-2.23 (m, 1H), 2.10-2.02 (m, 1H), 1.77-1.61 (m, 5H), 1.49-1.27 (m, 4H), 1.23-1.21 (m, 6H), 1.11-1.05 (m, 3H), 0.90-0.78 (m, 3H). $^{13}$C NMR (126 MHz, Methanol-d$_4$) δ 209.05, 176.25, 175.24, 173.92, 171.03, 169.14, 168.47, 159.32, 150.48, 148.89, 143.36, 138.87, 135.26, 131.02, 130.26 (2C), 129.44 (2C), 127.73, 121.74, 121.03, 115.66, 114.43, 113.63, 113.31, 78.06, 68.31, 56.59, 56.48, 55.56, 52.86, 50.88, 47.73, 45.74, 39.22, 38.42, 36.60, 36.19, 33.61, 32.28, 27.37, 25.93, 23.86, 23.66, 22.17, 17.40, 9.13. HRMS (ESI$^+$) m/z calcd for $C_{47}H_{61}N_5NaO_{11}^+$ [M+Na]$^+$: 894.4265, found 894.4261.

**SLF-F10:** *(R)-1-(3-(((4S,7S)-4-benzyl-7-methyl-3,6,9,13-tetraoxo-2-oxa-5,8,12-triazatetradecan-14-yl)oxy)phenyl)-3-(3,4-dimethoxyphenyl)propyl (S)-1-(3,3-dimethyl-2-oxopentanoyl)piperidine-2-carboxylate*

S-4 (45.0 mg, 0.062 mmol, 1.0 eq), HATU (30.8 mg, 0.081 mmol, 1.3 eq) and L-Phenylalanine methyl ester hydrochloride (26.7 mg, 0.124 mmol, 2.0 eq) were dissolved in dry DMF (2.0 mL) at room temperature. DIPEA (51 µL, 0.310 mmol, 5.0 eq) was added to the reaction and stirred for 1 h. Upon completion, the reaction was concentrated, extracted with DCM (20 mL) and washed with brine (1 x 20 mL). The organic layer was dried by $Na_2SO_4$, concentrated, and the resulting residue purified by flash chromatography to provide SLF-F10 as white powder (37.0 mg, 0.042 mmol, 67.7%).

$^1H$ NMR (500 MHz, $CDCl_3$) δ 7.37-7.34 (m, 1H), 7.26-7.17 (m, 4H), 7.09-7.07 (m, 2H), 6.97-6.93 (m, 1H), 6.92-6.89 (m, 1H), 6.83-6.81 (m, 1H), 6.78-6.76 (m, 1H), 6.69-6.64 (m, 3H), 6.49-6.46 (m, 1H), 5.76-5.73 (m, 1H), 5.29 (s, 1H), 4.81-4.76 (m, 1H), 4.46-4.45 (m, 2H), 4.37-4.35 (m, 1H), 3.84 (s, 3H), 3.83 (s, 3H), 3.67 (s, 3H), 3.62-3.55 (m, 2H), 3.38-3.35 (m, 1H), 3.21-3.10 (m, 2H), 3.04-3.00 (m, 1H), 2.62-2.50 (m, 2H), 2.44-2.41 (m, 2H), 2.39-2.36 (m, 1H), 2.26-2.19 (m, 1H), 2.07-2.00 (m, 1H), 1.79-1.61 (m, 5H), 1.36-1.25 (m, 5H), 1.21-1.11 (m, 6H), 0.88-0.78 (m, 3H). $^{13}C$ NMR (126 MHz, $CDCl_3$) δ 207.91, 171.85, 171.73, 171.19, 169.67, 168.52, 167.35, 157.42, 148.89, 147.37, 141.81, 135.81, 133.35, 129.97, 129.22 (2C), 128.55 (2C), 127.11, 120.16, 120.06, 114.17, 113.28, 111.74, 111.35, 76.54, 67.22, 55.91, 55.83, 53.31, 52.35, 51.33, 48.96, 46.72, 44.19, 38.16, 37.67, 32.45, 31.25, 26.44, 24.92, 23.46, 23.12, 21.19, 18.53, 17.94, 17.16, 8.75. HRMS ($ESI^+$) m/z calcd for $C_{48}H_{62}N_4NaO_{12}^+$ $[M+Na]^+$: 909.4262, found 909.4259.

**SLF-F11:** *(3-(2-(3-((R)-3-(3,4-dimethoxyphenyl)-1-(((S)-1-(3,3-dimethyl-2-oxopentanoyl)piperidine-2-carbonyl)oxy)propyl)phenoxy)acetamido)propanoyl)-L-alanyl-L-phenylalanine*

SLF-F10 (20.5 mg, 0.023 mmol, 1.0 eq) was dissolved in MeOH/$H_2O$ (2.0 mL, 4:1), $LiOH\cdot H_2O$(9.66 mg, 0.230 mmol, 10 eq) was added to the reaction and stirred at 0℃ for 0.5 h. 1M HCl was added to the reaction and pH was adjusted to 2, then concentrated and purified by flash chromatography to provide SLF-F11 as white powder (9.8 mg, 0.011 mmol, 47.8%).

$^1H$ NMR (500 MHz, Methanol-$d_4$) δ 7.32-7.28 (m, 1H), 7.24-7.19 (m, 4H), 7.17-7.14 (m, 1H), 7.01-6.92 (m, 3H), 6.86-6.84 (m, 1H), 6.80-6.77 (m, 1H), 6.72-6.70 (m, 1H), 5.77-5.73 (m, 1H), 5.23-5.22 (m, 1H), 4.60 (s, 1H), 4.50 (s, 2H), 4.33-4.29 (m, 1H), 3.80 (s, 3H), 3.79 (s, 3H), 3.75-3.70 (m, 1H), 3.53 (t, *J* = 7.0 Hz, 2H), 3.42-3.38 (m, 1H), 3.23-3.16 (m, 2H), 3.03-2.99 (m, 1H), 2.66-2.54 (m, 2H), 2.45 (t, *J* = 6.5 Hz, 2H), 2.36-2.33 (m, 1H),

2.29-2.21 (m, 1H), 2.09-2.02 (m, 1H), 1.76-1.61 (m, 5H), 1.32-1.26 (m, 4H), 1.22-1.10 (m, 6H), 0.89-0.78 (m, 3H). $^{13}C$ NMR (126 MHz, Methanol-$d_4$) δ 209.04, 174.67, 173.65, 171.01, 169.14, 168.47, 159.30, 150.47, 148.88, 143.37, 143.16, 138.47, 135.26, 131.02 (2C), 130.51 (2C), 129.40, 127.75, 121.73, 121.05, 115.63, 114.47, 113.63, 113.30, 78.02, 68.29, 56.59, 56.48, 55.88, 52.85, 50.45, 47.73, 45.74, 39.23, 38.37, 36.60, 36.30, 33.60, 32.28, 27.37, 25.92, 23.87, 23.66, 22.16, 17.85, 9.13. HRMS (ESI$^+$) m/z calcd for $C_{47}H_{60}N_4NaO_{12}^+$ [M+Na]$^+$: 895.4105, found 895.4103.

**Synthesis of MKI-F3 and MKI-F4**

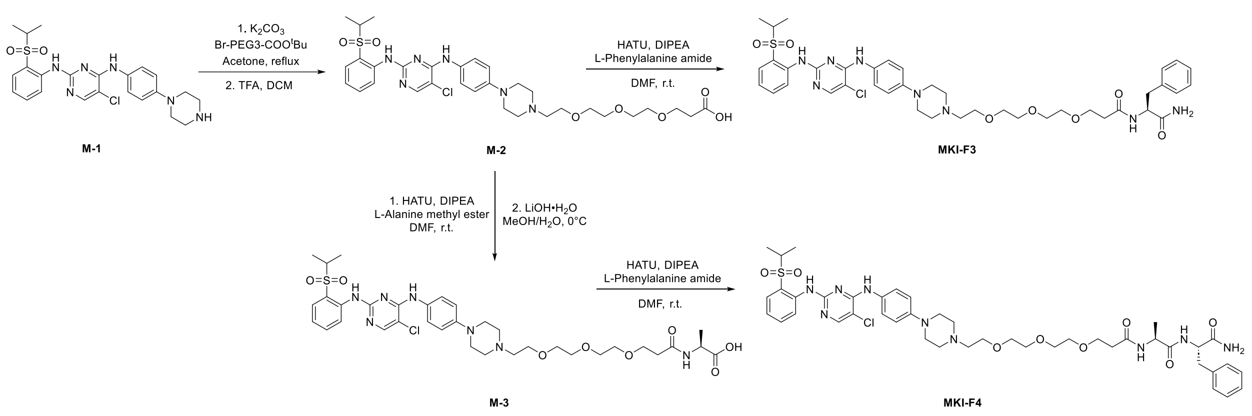


**MKI-F3:** *(S)-N-(1-amino-1-oxo-3-phenylpropan-2-yl)-3-(2-(2-(2-(4-(4-((5-chloro-2-((2-(isopropylsulfonyl)phenyl)amino)pyrimidin-4-yl)amino)phenyl)piperazin-1-yl)ethoxy)ethoxy)ethoxy)propenamide*

Step1: M-1 (100 mg, 0.205 mmol, 1.0 eq), $K_2CO_3$ (56.6 mg, 0.41 mmol, 2.0 eq) and Br-PEG3-COO$^t$Bu (83.9 mg, 0.246 mmol, 1.2 eq) were dissolved in acetone and refluxed overnight. Upon completion, the reaction was concentrated, extracted with EtOAc (30 mL) and washed with brine (1 x 20 mL). The organic layer was dried by $Na_2SO_4$, concentrated, and the resulting residue was used without further purification. The crude product was dissolved in DCM (2.0 mL), TFA (0.5 mL) was added to the reaction and stirred for 2 h. The reaction mixture was then concentrated and purified by flash chromatography to provide M-2 as light yellow powder (76.6 mg, 0.111 mmol, 54.1% over 2 steps).

$^1H$ NMR (500 MHz, Methanol-$d_4$) δ 8.65 (d, *J* = 8.5 Hz, 1H), 8.12 (s, 1H), 7.89 (dd, *J* = 8.0 Hz, 1.5 Hz, 1H), 7.67 (t, *J* = 8.0 Hz, 1H), 7.47 (d, *J* = 9.0 Hz, 2H), 7.34-7.31 (m, 1H), 6.99 (d, *J* = 9.0 Hz, 2H), 3.89 (t, *J* = 5.0 Hz, 2H), 3.74 (t, *J* = 6.0 Hz, 2H), 3.71-3.68 (m, 4H), 3.66-3.61 (m, 4H), 3.46-3.42 (m, 8H), 3.38-3.33 (m, 3H), 2.52 (t, *J* = 6.0 Hz, 1H), 1.26 (d, *J* = 6.5 Hz, 6H).

Step2: M-2 (36.6 mg, 0.053 mmol, 1.0 eq), HATU (26.2 mg, 0.069 mmol, 1.3 eq) and L-Phenylalanine amide (17.4 mg, 0.106 mmol, 2.0 eq) were dissolved in dry DMF (2.0 mL) at room temperature. DIPEA (44 μL, 0.265 mmol, 5.0 eq) was added to the reaction and stirred for 1 h. Upon completion, the reaction was concentrated, extracted with DCM (20 mL) and washed with brine (1 x 20 mL). The organic layer was dried by $Na_2SO_4$, concentrated, and the resulting residue purified by flash chromatography to provide MKI-F3 as light yellow powder (19.8 mg, 0.024 mmol, 45.3%).

$^1H$ NMR (500 MHz, Methanol-$d_4$) δ 8.55-8.53 (m, 1H), 8.11 (s, 1H), 7.89 (d, *J* = 8.0 Hz, 1H), 7.67 (t, *J* = 7.5 Hz, 1H), 7.43 (d, *J* = 8.5 Hz, 1H), 7.37-7.34 (m, 1H), 7.27-7.23 (m, 4H), 7.20-7.17 (m, 1H), 6.99 (d, *J* = 8.0 Hz, 1H), 4.61-4.58 (m, 1H), 3.89 (t, *J* = 4.5 Hz, 1H), 3.78-3.70 (m, 8H), 3.65-3.58 (m, 8H), 3.49-3.47 (m, 2H), 3.24-3.20 (m, 2H), 3.11-3.03 (m, 2H), 2.90-2.85 (m, 2H), 2.52-2.46 (m, 1H), 2.37-2.32 (m, 1H), 1.24 (d, *J* = 6.5 Hz, 6H). $^{13}C$ NMR (126 MHz, Methanol-$d_4$) δ 176.48, 174.11, 158.27, 157.39, 153.09, 147.70, 139.17, 138.79, 136.03, 133.97, 132.37, 130.21, 129.50, 127.80, 126.76, 125.74, 125.32, 124.05, 118.52, 106.43, 71.72, 71.55, 71.46, 71.21, 68.60, 65.38, 57.17, 56.94, 55.94, 55.87, 53.36, 38.48, 37.60, 15.49. HRMS (ESI$^+$) m/z calcd for $C_{41}H_{53}ClN_8NaO_7S^+$ $[M+Na]^+$: 859.3344, found 859.3372.

**MKI-F4:** *(S)-N-((S)-1-amino-1-oxo-3-phenylpropan-2-yl)-2-(3-(2-(2-(2-(4-(4-((5-chloro-2-((2-(isopropylsulfonyl)phenyl)amino)pyrimidin-4-yl)amino)phenyl)piperazin-1-yl)ethoxy)ethoxy)ethoxy)propanamido)propenamide*

Step1: M-2 (40.0 mg, 0.058 mmol, 1.0 eq), HATU (28.5 mg, 0.075 mmol, 1.3 eq) and L-Alanine methyl ester hydrochloride (36.3 mg, 0.260 mmol, 2.0 eq) were dissolved in dry

DMF (3.0 mL) at room temperature. DIPEA (48 μL, 0.290 mmol, 5.0 eq) was added to the reaction and stirred for 1 h. Upon completion, the reaction was concentrated, extracted with DCM (30 mL) and washed with brine (1 x 30 mL). The organic layer was dried by $Na_2SO_4$, concentrated, and the resulting residue was used without further purification. The crude product was dissolved in MeOH/$H_2O$ (3.0 mL, 4:1), LiOH·$H_2O$(24.4 mg, 0.580 mmol, 10 eq) was added to the reaction and stirred at 0℃ for 0.5 h. 1M HCl was added to the reaction and pH was adjusted to 2, then concentrated and purified by flash chromatography to provide M-3 as light yellow powder (26.2 mg, 0.034 mmol, 58.6% over 2 steps).

$^1$H NMR (500 MHz, Methanol-d$_4$) δ 8.65 (d, *J* = 8.5 Hz, 1H), 8.11 (s, 1H), 7.89-7.87 (m, 1H), 7.66 (t, *J* = 8.0 Hz, 1H), 7.47 (d, *J* = 8.5 Hz, 2H), 7.32 (t, *J* = 7.5 Hz, 1H), 6.99-6.97 (m, 2H), 4.32 (s, 1H), 3.88 (t, *J* = 5.0 Hz, 2H), 3.77-3.73 (m, 2H), 3.69-3.62 (m, 8H), 3.50-3.48 (m, 4H), 3.43-3.39 (m, 6H), 3.35 (s, 1H), 2.50-2.47 (m, 2H), 1.37 (d, *J* = 7.0 Hz, 3H), 1.25 (d, *J* = 7.0 Hz, 6H).

Step2: M-3 (26.2 mg, 0.034 mmol, 1.0 eq), HATU (16.7 mg, 0.044 mmol, 1.3 eq) and L-Phenylalanine amide (11.2 mg, 0.068 mmol, 2.0 eq) were dissolved in dry DMF (2.0 mL) at room temperature. DIPEA (28 μL, 0.170 mmol, 5.0 eq) was added to the reaction and stirred for 1 h. Upon completion, the reaction was concentrated, extracted with DCM (20 mL) and washed with brine (1 x 20 mL). The organic layer was dried by $Na_2SO_4$, concentrated, and the resulting residue purified by flash chromatography to provide MKI-F3 as light yellow powder (11.0 mg, 0.012 mmol, 35.3%).

$^1$H NMR (500 MHz, Methanol-d$_4$) δ 8.64 (d, *J* = 8.5 Hz, 1H), 8.09 (s, 1H), 7.88-7.86 (m, 1H), 7.67-7.63 (m, 1H), 7.47 (d, *J* = 9.0 Hz, 2H), 7.33-7.29 (m, 1H), 7.27-7.20 (m, 4H), 7.20-7.16 (m, 1H), 6.97 (d, *J* = 8.5 Hz, 2H), 4.59-4.56 (m, 1H), 4.20 (q, *J* = 7.0 Hz, 1H), 3.88 (t, *J* = 5.0 Hz, 2H), 3.76-3.60 (m, 12H), 3.45-3.43 (m, 2H), 3.33-3.27 (m, 4H), 3.26-3.18 (m, 2H), 3.08-3.03 (m, 1H), 2.95-2.91 (m, 1H), 2.86-2.82 (m, 1H), 2.48 (t, *J* = 6.0 Hz, 2H), 1.24 (d, *J* = 6.0 Hz, 6H), 1.20 (d, *J* = 7.5 Hz, 3H). $^{13}$C NMR (126 MHz, Methanol-d$_4$) δ 175.94, 174.93, 174.16, 159.61, 156.74, 155.95, 147.02, 139.76, 138.56, 135.95,

132.28, 130.50, 130.36 (2C), 130.13, 129.48 (2C), 127.80, 125.89, 125.16, 124.51, 123.29, 118.57, 106.23, 71.39-71.36 (6C), 68.03, 65.31, 57.32, 56.92, 55.51, 53.46, 51.12, 49.88, 38.52, 37.15, 17.60, 15.52 (2C). HRMS (ESI$^+$) m/z calcd for $C_{44}H_{59}ClN_9O_8S^+$ [M+H]$^+$: 908.3896, found 908.3924.

**Synthesis of JQ1-F4**

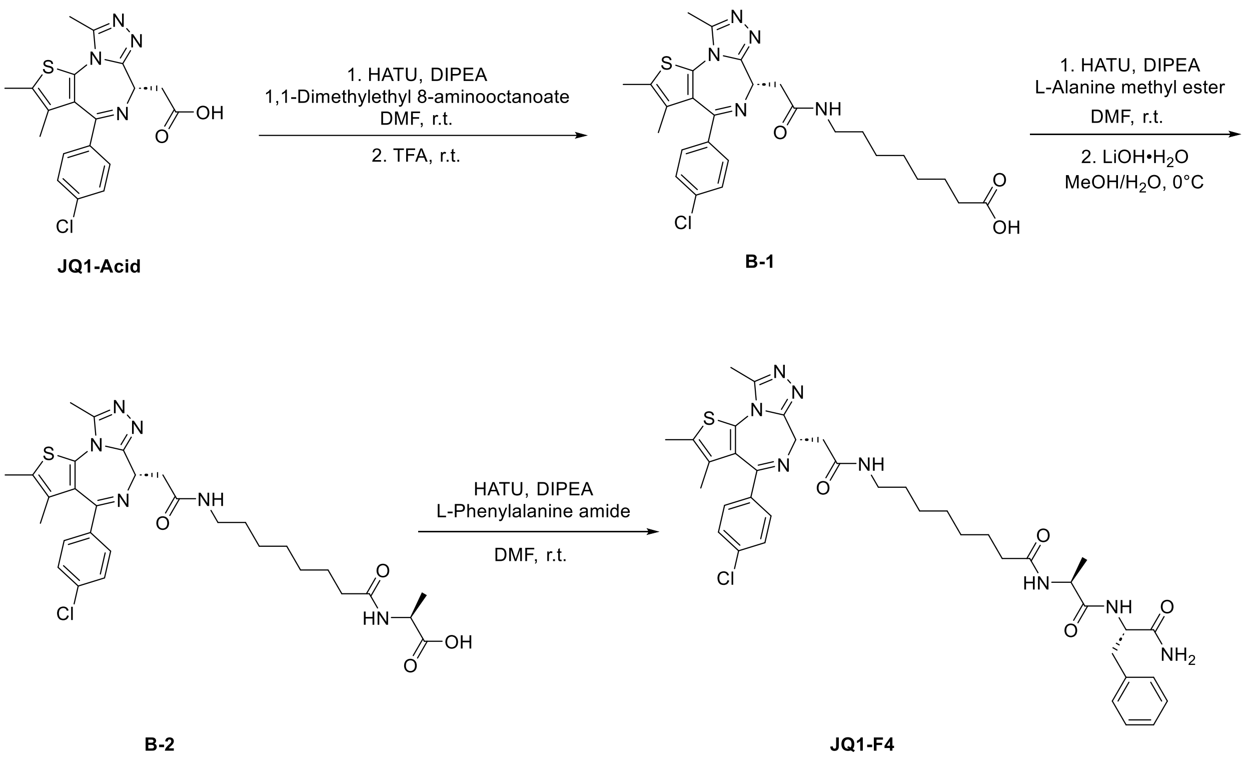


**JQ-F4:** *N-((S)-1-(((S)-1-amino-1-oxo-3-phenylpropan-2-yl)amino)-1-oxopropan-2-yl)-8-(2-((S)-4-(4-chlorophenyl)-2,3,9-trimethyl-6H-thieno[3,2-f][1,2,4]triazolo[4,3-a][1,4]diazepin-6-yl)acetamido)octanamide*

Step1: JQ-1 Acid (100 mg, 0.25 mmol, 1.0 eq), HATU (124 mg, 0.325 mmol, 1.3 eq) and 1,1-dimethylethyl 8-aminooctanoate (108 mg, 0.50 mmol, 2.0 eq) were dissolved in dry DMF (3.0 mL) at room temperature. DIPEA (207 µL, 1.25 mmol, 5.0 eq) was added to the reaction and stirred for 1 h. Upon completion, the reaction was concentrated, extracted with DCM (30 mL) and washed with brine (1 x 30 mL). The organic layer was dried by

$Na_2SO_4$, concentrated, and the resulting residue was used without further purification. The crude product was dissolved in DCM (5.0 mL), TFA (1.0 mL) was added to the reaction and stirred for 2 h. The reaction mixture was then concentrated and purified by flash chromatography to provide B-1 as white powder (56.0 mg, 0.103 mmol, 41.2% over 2 steps).

$^1$H NMR (500 MHz, $CDCl_3$) δ 7.43-7.38 (m, 4H), 4.87 (t, $J$ = 7.0 Hz, 1H), 3.65-3.60 (m, 1H), 3.53-3.49 (m, 1H), 3.33-3.28 (m, 2H), 2.82 (s, 3H), 2.45 (s, 3H), 2.36 (t, $J$ = 7.0 Hz, 2H), 1.69 (s, 3H), 1.64-1.60 (m, 2H), 1.57-1.54 (m, 2H), 1.36-1.33 (m, 6H).

Step2: B-1 (56.0 mg, 0.103 mmol, 1.0 eq), HATU (51.0 mg, 0.134 mmol, 1.3 eq) and L-Alanine methyl ester hydrochloride (28.8 mg, 0.206 mmol, 2.0 eq) were dissolved in dry DMF (3.0 mL) at room temperature. DIPEA (85 μL, 0.515 mmol, 5.0 eq) was added to the reaction and stirred for 1 h. Upon completion, the reaction was concentrated, extracted with DCM (30 mL) and washed with brine (1 x 30 mL). The organic layer was dried by $Na_2SO_4$, concentrated, and the resulting residue was used without further purification. The crude product was dissolved in MeOH/$H_2O$ (3.0 mL, 4:1), LiOH·$H_2O$(43.3 mg, 1.03 mmol, 10 eq) was added to the reaction and stirred at 0℃ for 0.5 h. 1M HCl was added to the reaction and pH was adjusted to 2, then concentrated and purified by flash chromatography to provide B-2 as white powder (35.0 mg, 0.057 mmol, 55.3% over 2 steps).

$^1$H NMR (500 MHz, Methanol-$d_4$) δ 7.48 (d, $J$ = 8.5 Hz, 2H), 7.43 (d, $J$ = 8.5 Hz, 2H), 4.74-4.71 (m, 1H), 4.37 (q, $J$ = 7.5 Hz, 1H), 3.46-3.41 (m, 1H), 3.35-3.34 (m, 1H), 3.27-3.25 (m, 2H), 2.77 (s, 3H), 2.47 (s, 3H), 2.24-2.21 (m, 2H), 1.71 (s, 3H), 1.63-1.55 (m, 4H), 1.38-1.37 (m, 11H).

Step3: B-2 (35.0 mg, 0.057 mmol, 1.0 eq), HATU (28.2 mg, 0.074 mmol, 1.3 eq) and L-Phenylalanine amide (18.7 mg, 0.114 mmol, 2.0 eq) were dissolved in dry DMF (2.0 mL) at room temperature. DIPEA (47 μL, 0.285 mmol, 5.0 eq) was added to the reaction and stirred for 1 h. Upon completion, the reaction was concentrated, extracted with DCM (20 mL) and washed with brine (1 x 20 mL). The organic layer was dried by $Na_2SO_4$, concentrated, and the resulting residue purified by flash chromatography to provide JQ1-F4 as white powder (16.2 mg, 0.021 mmol, 36.8%).

$^{1}$H NMR (500 MHz, Methanol-$d_4$) δ 7.47-7.45 (d, *J* = 7.0 Hz, 2H), 7.42-7.40 (d, *J* = 7.0 Hz, 2H), 7.27-7.16 (m, 5H), 4.69-4.66 (m, 1H), 4.59-4.56 (m, 1H), 4.15-4.11 (m, 1H), 3.44-3.40 (m, 1H), 3.33-3.29 (m, 1H), 3.27-3.24 (m, 2H), 3.18-3.14 (m, 1H), 2.99-2.94 (m, 1H), 2.72 (s, 3H), 2.45 (s, 3H), 2.19 (t, *J* = 7.5 Hz, 2H), 1.70 (s, 3H), 1.61-1.54 (m, 4H), 1.40-1.29 (m, 6H), 1.22 (d, *J* = 7.5 Hz, 3H). $^{13}$C NMR (126 MHz, Methanol-$d_4$) δ 176.48, 175.86, 174.90, 172.49, 166.46, 156.96, 152.39, 138.43, 138.24, 137.81, 133.65, 133.46, 132.17, 132.04, 131.44 (2C), 130.40 (2C), 129.84 (2C), 129.45 (2C), 127.77, 55.45, 55.11, 50.78, 40.49, 38.61, 38.59, 36.65, 30.42, 30.23, 30.08, 27.83, 26.65, 17.58, 14.44, 12.98, 11.60. HRMS (ESI$^+$) m/z calcd for $C_{39}H_{47}ClN_8NaO_4S^+$ [M+Na]$^+$: 781.3027, found 781.3019.

### $^1$H NMR spectrum of SLF-F1

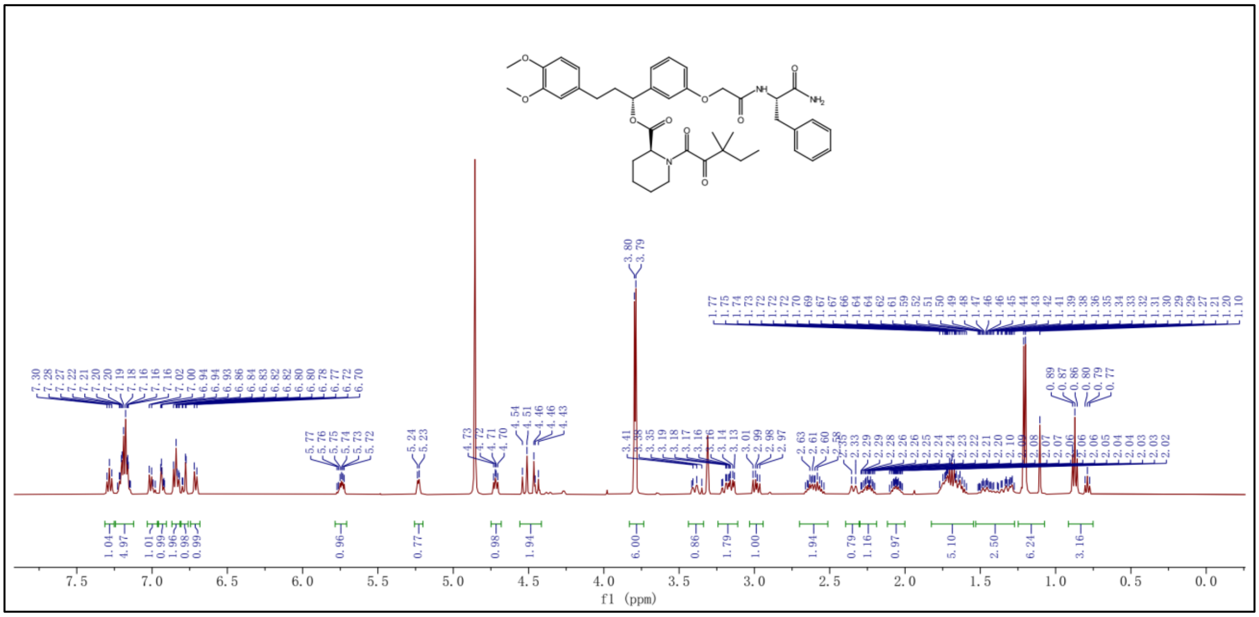

### $^{13}$C NMR spectrum of SLF-F1

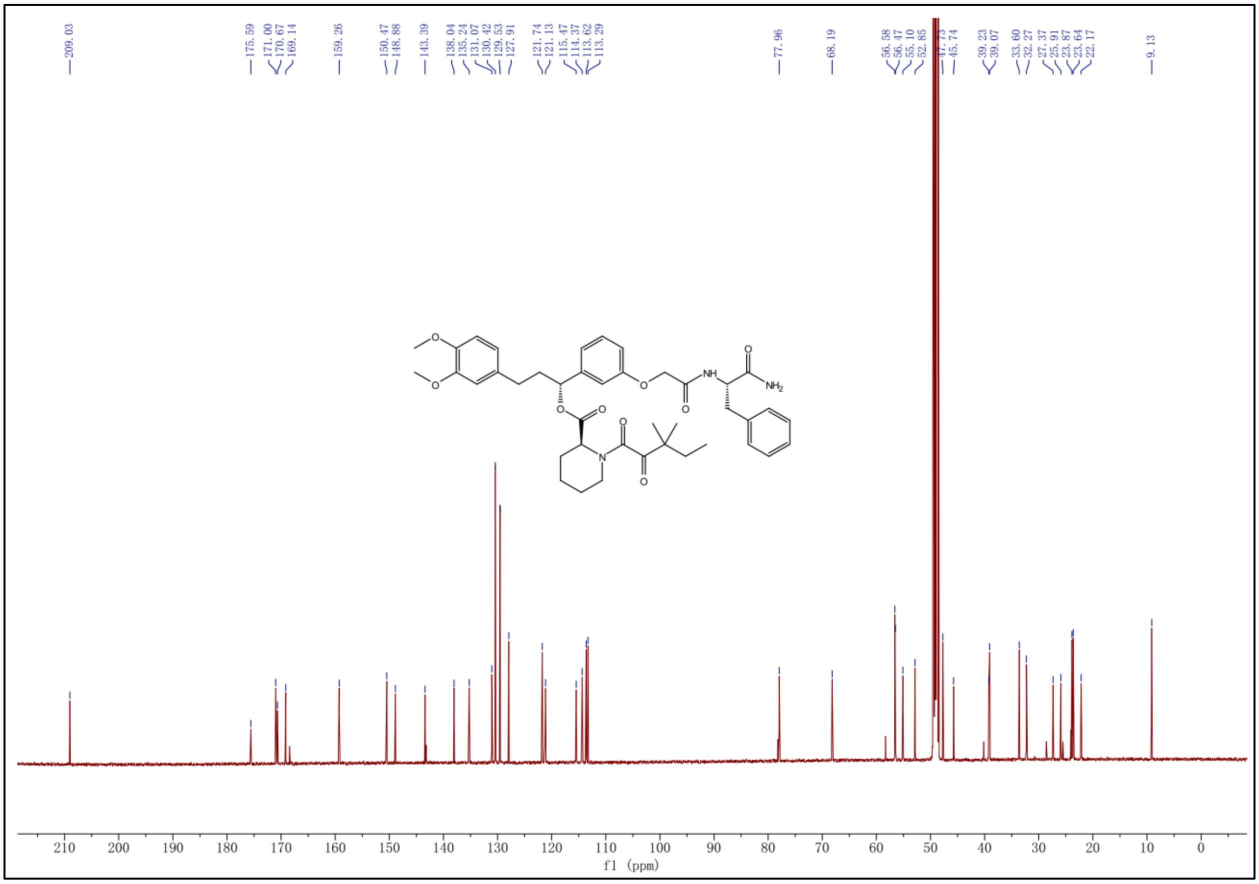

**[1]H NMR spectrum of SLF-F2**

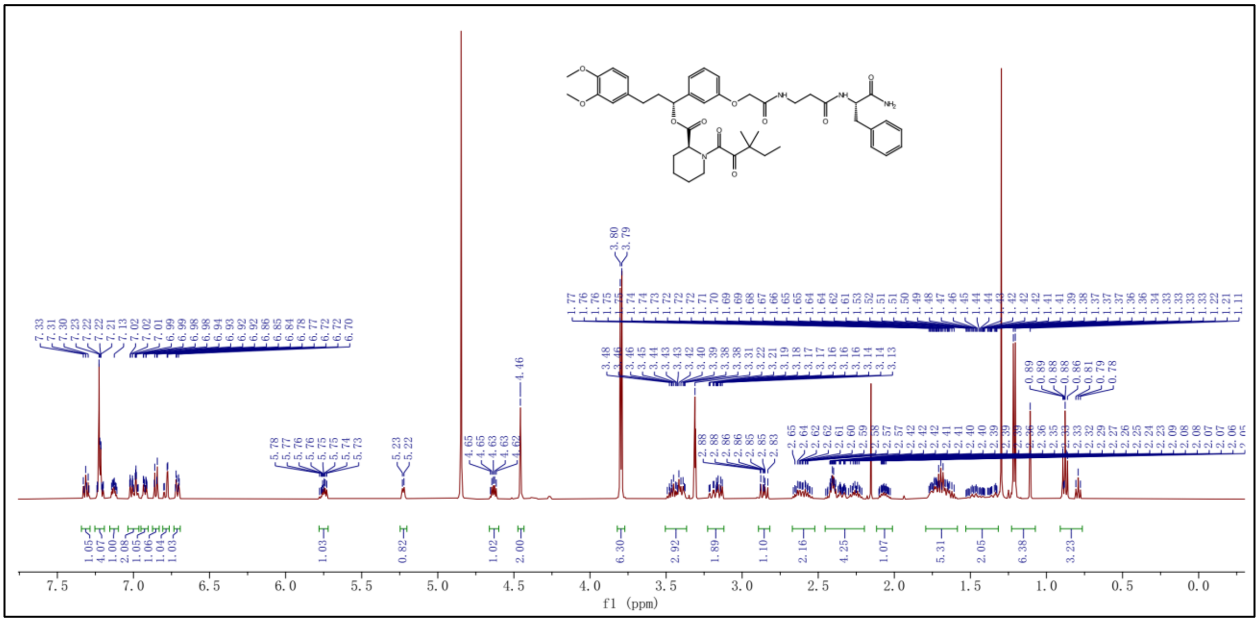

**[13]C NMR spectrum of SLF-F2**

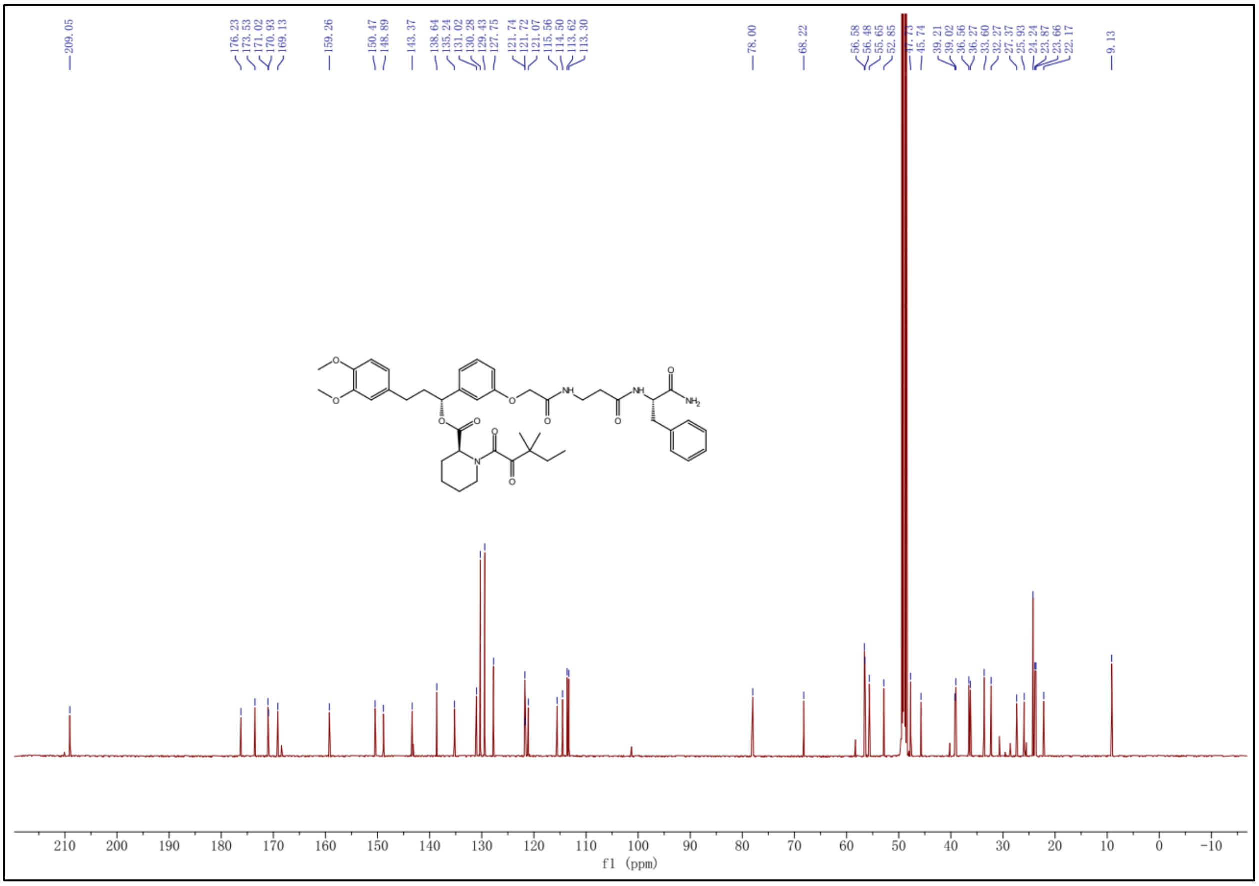

[1]H NMR spectrum of SLF-F3

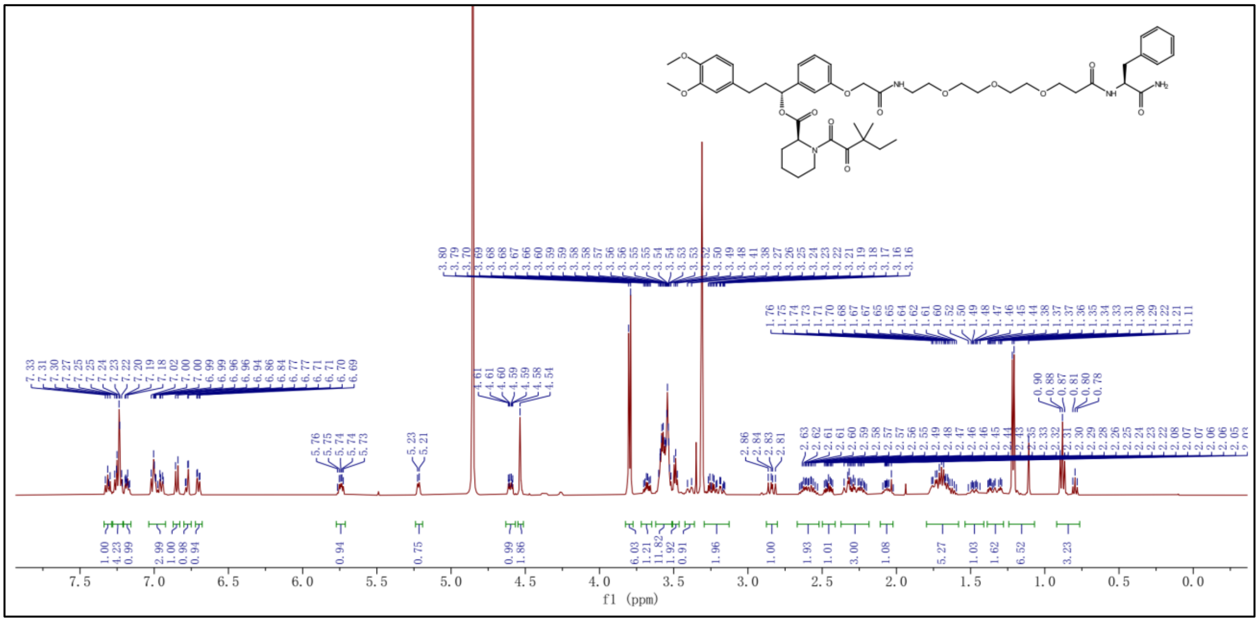

[13]C NMR spectrum of SLF-F3

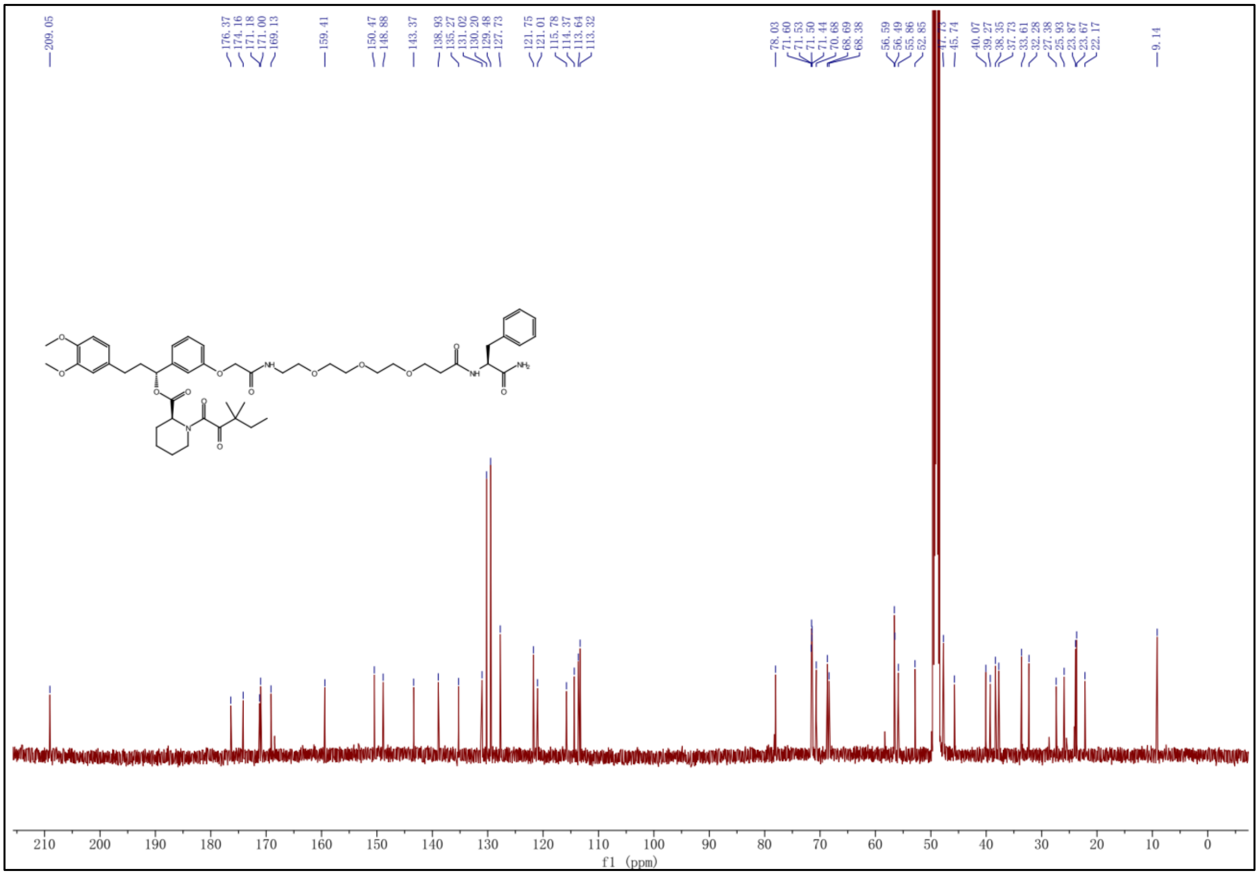

#### $^{1}$H NMR spectrum of SLF-F4

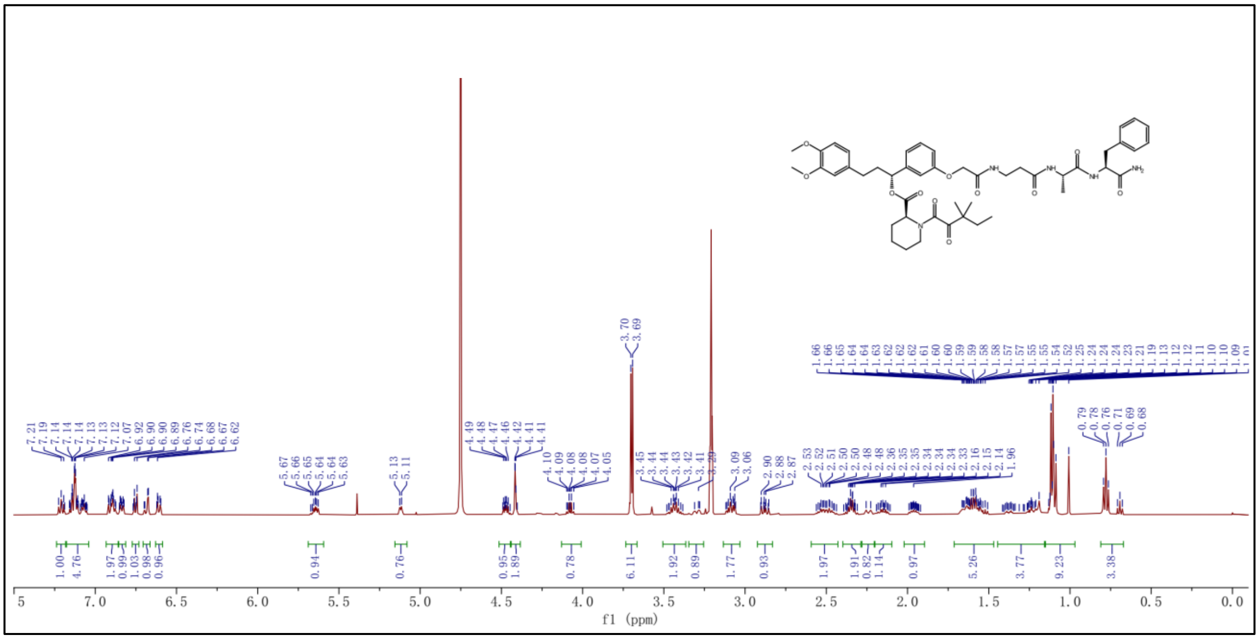

#### $^{13}$C NMR spectrum of SLF-F4

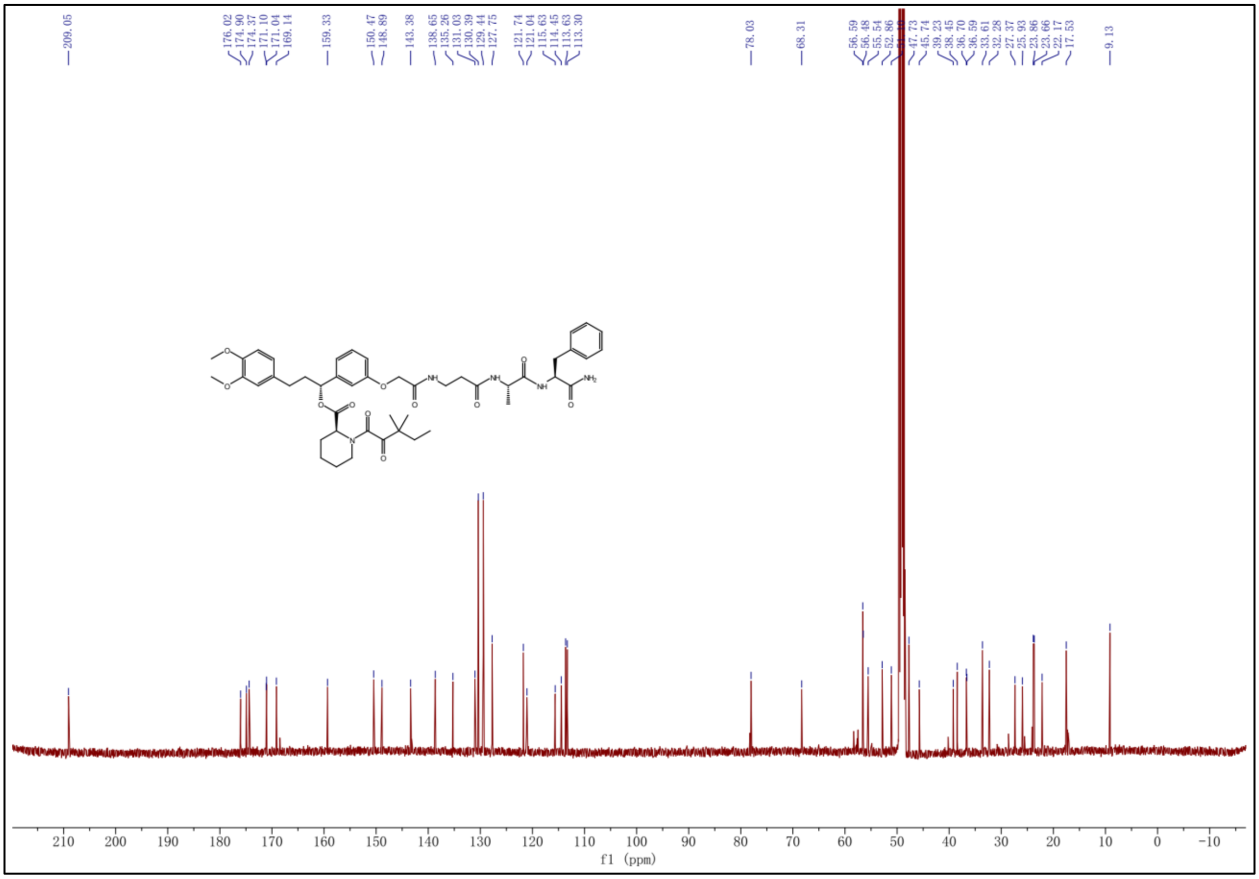

### $^{1}$H NMR spectrum of SLF-F5

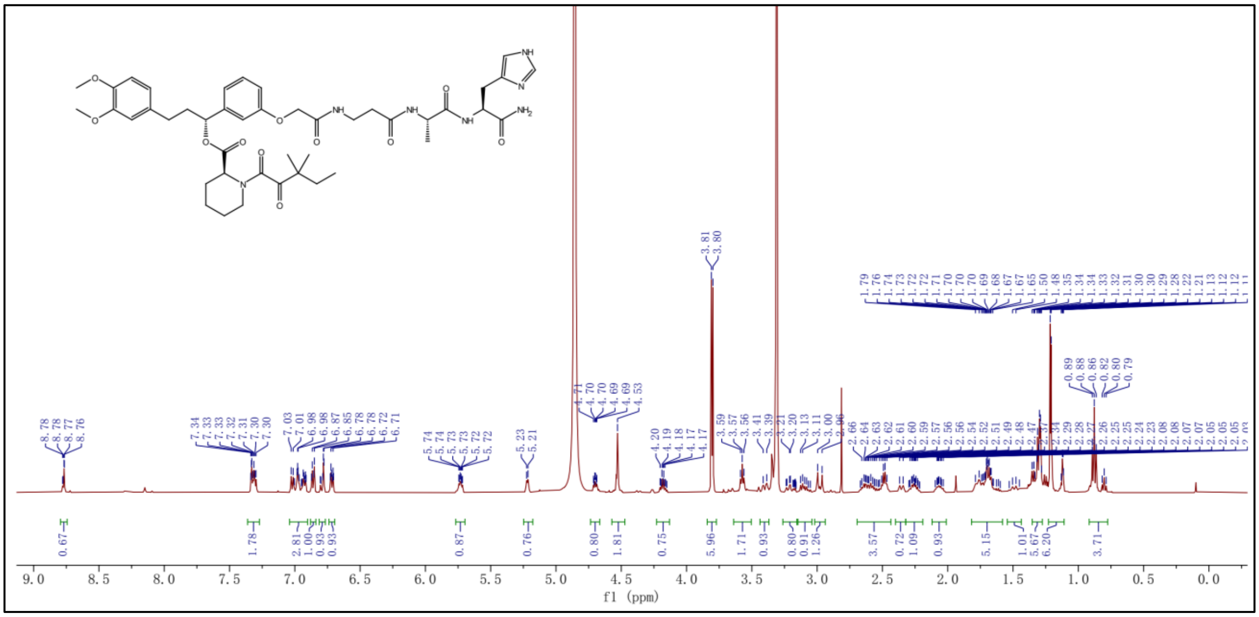

### $^{13}$C NMR spectrum of SLF-F5

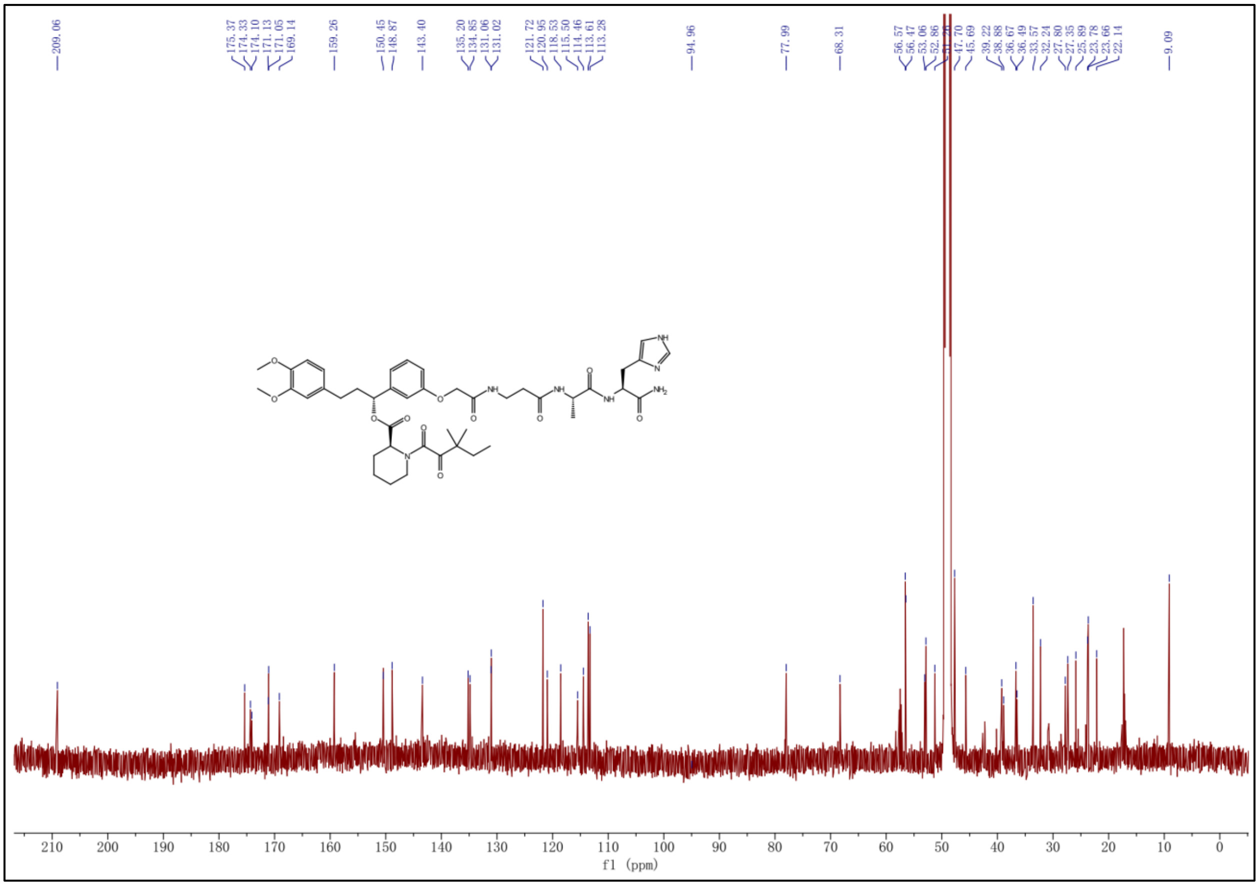

<sup></sup>

### $^{1}$H NMR spectrum of SLF-F6

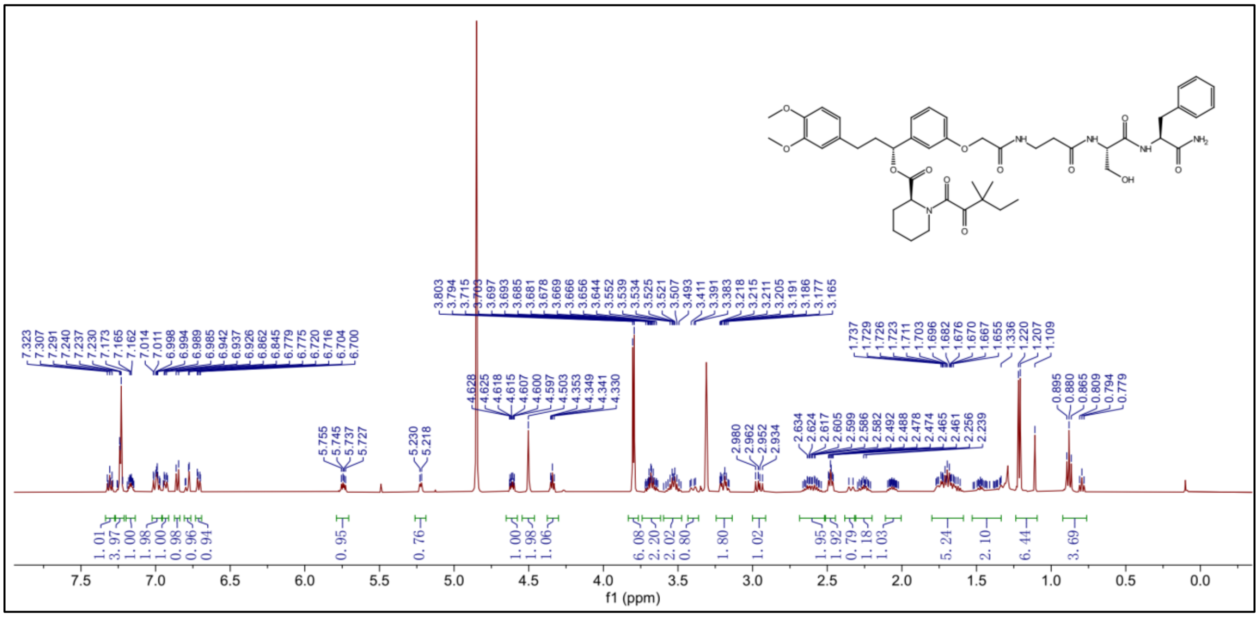

### $^{13}$C NMR spectrum of SLF-F6

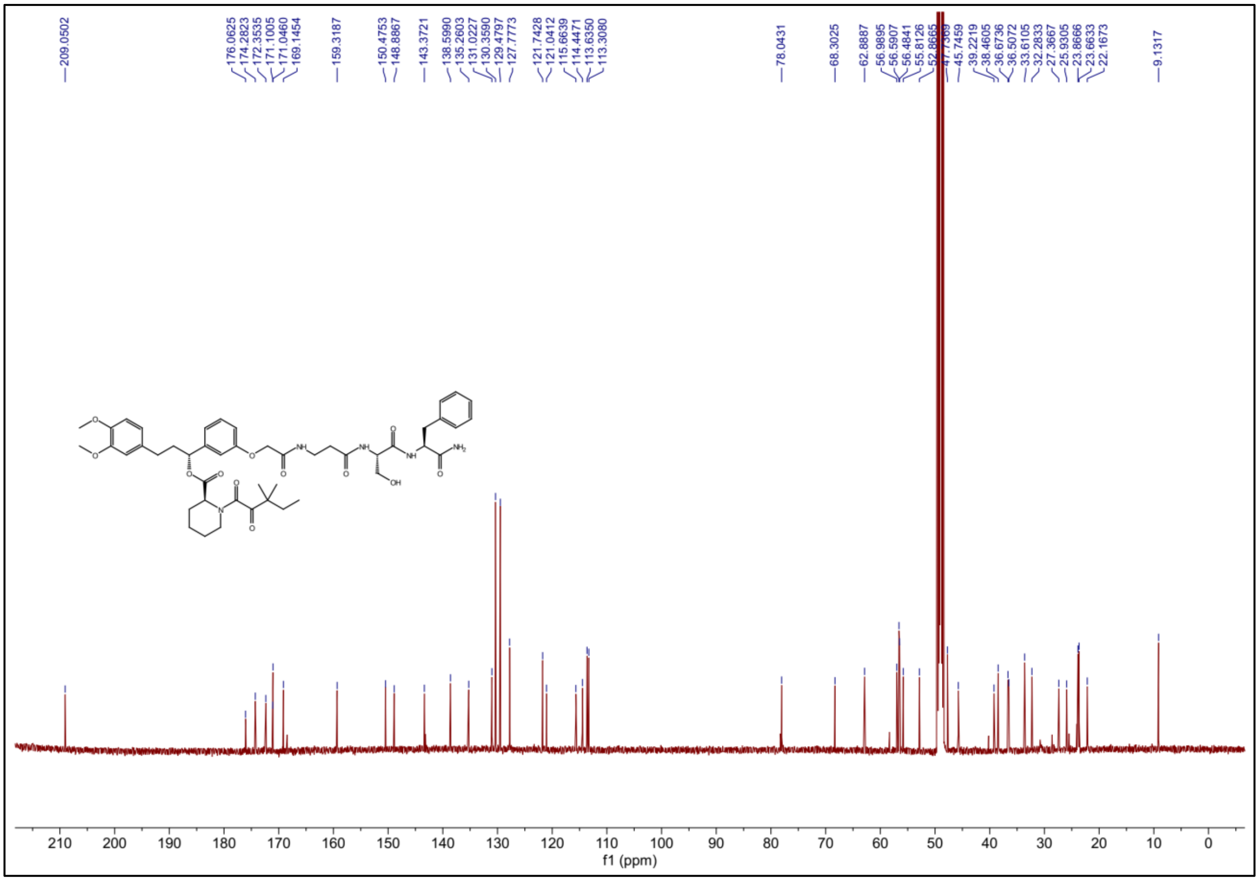

### [1]H NMR spectrum of SLF-F7

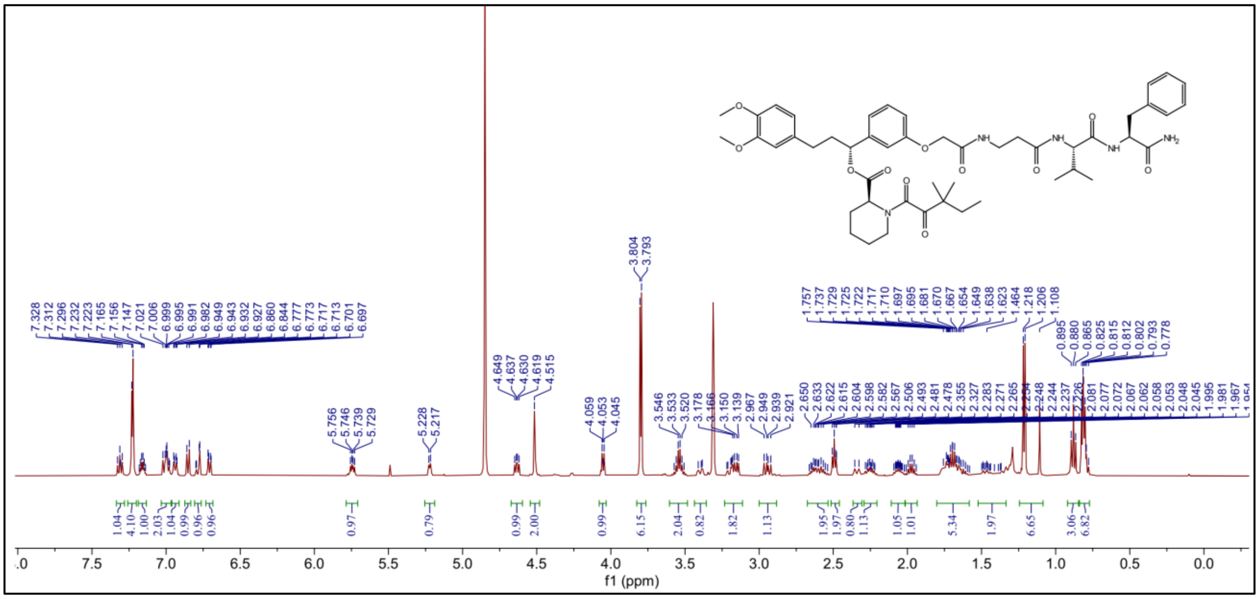

### [13]C NMR spectrum of SLF-F7

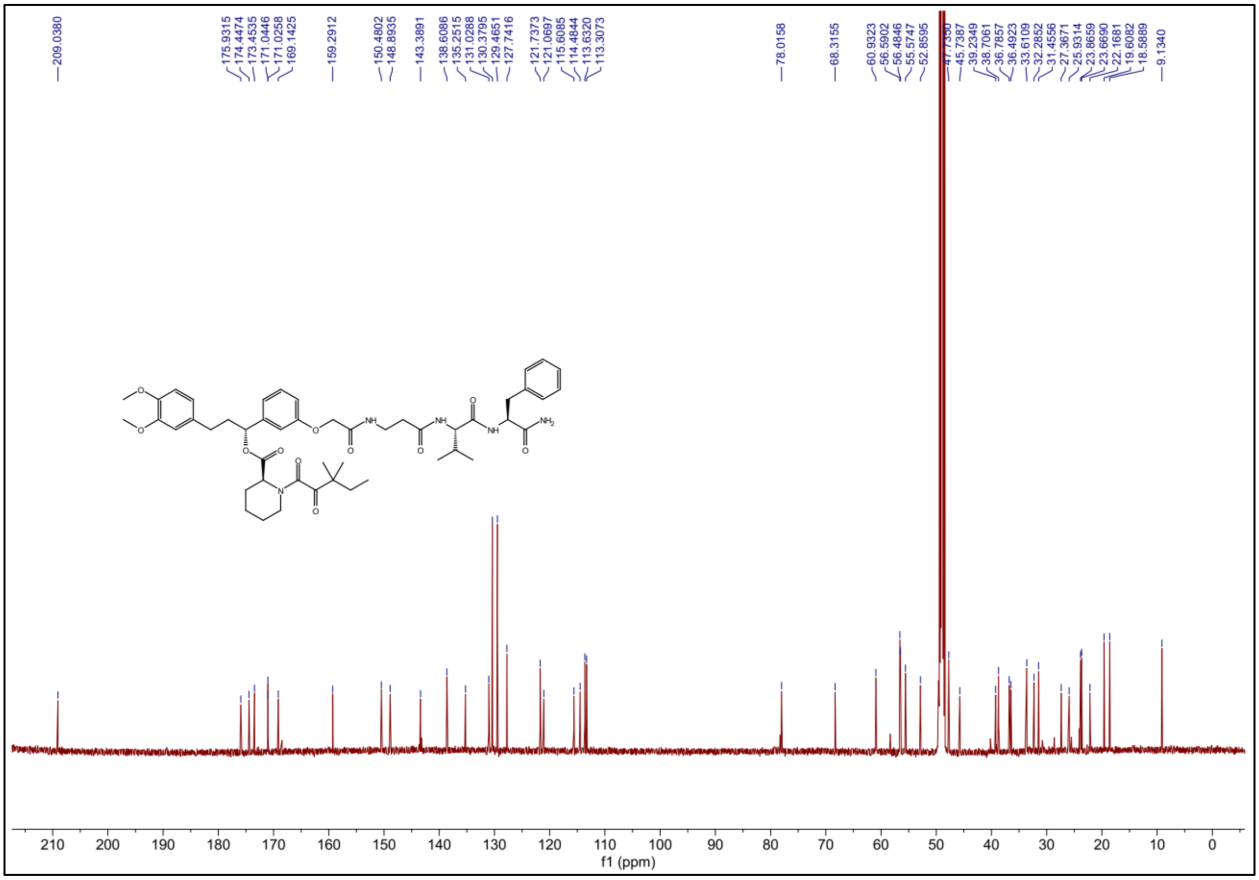

### $^{1}$H NMR spectrum of SLF-F8

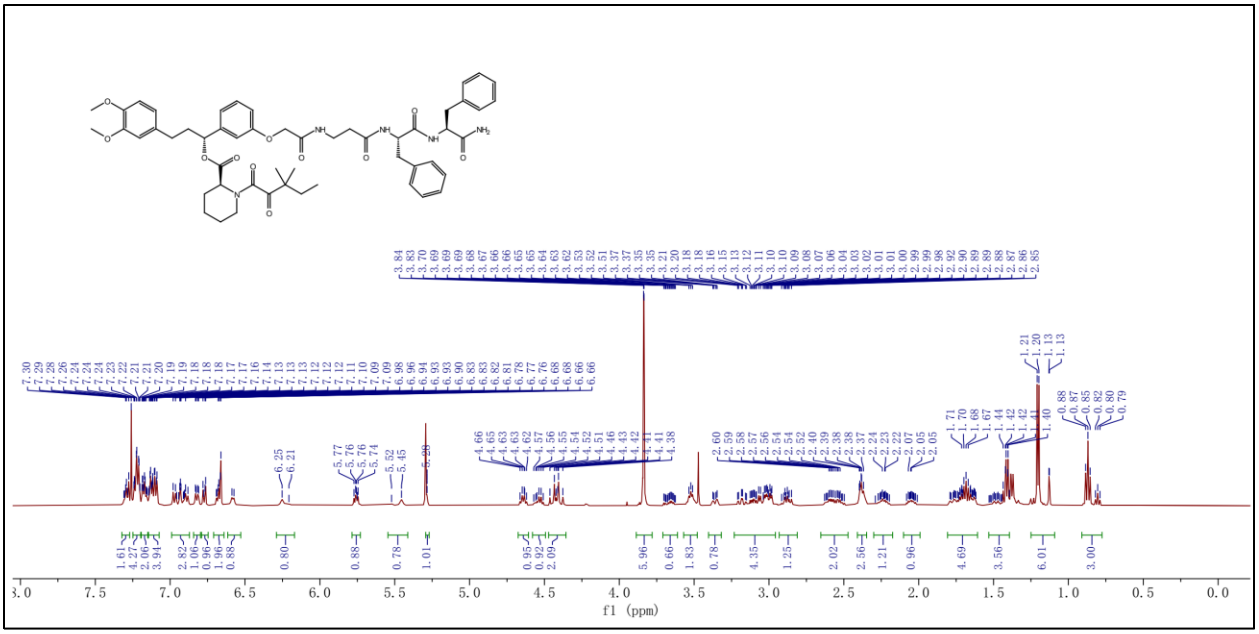

### $^{13}$C NMR spectrum of SLF-F8

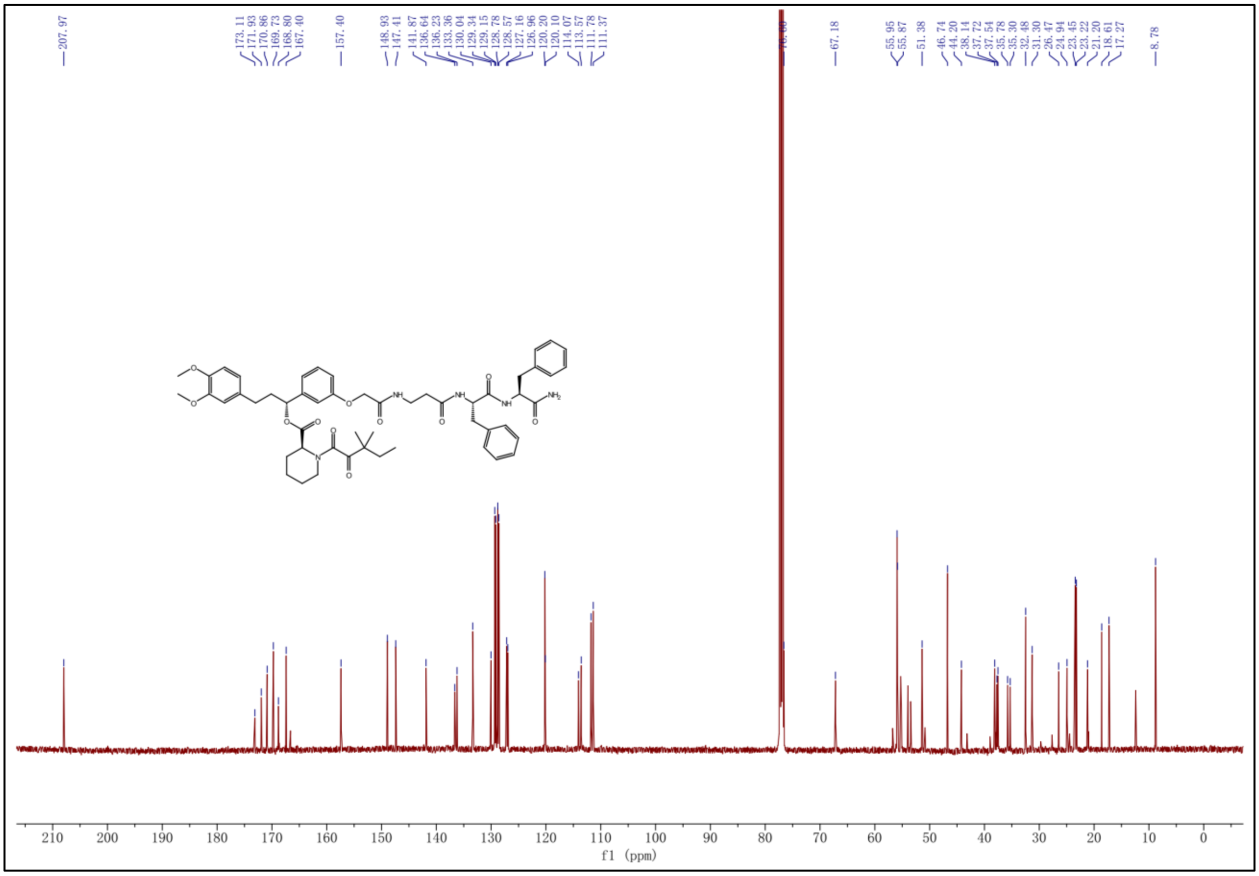

### $^1$H NMR spectrum of SLF-F9

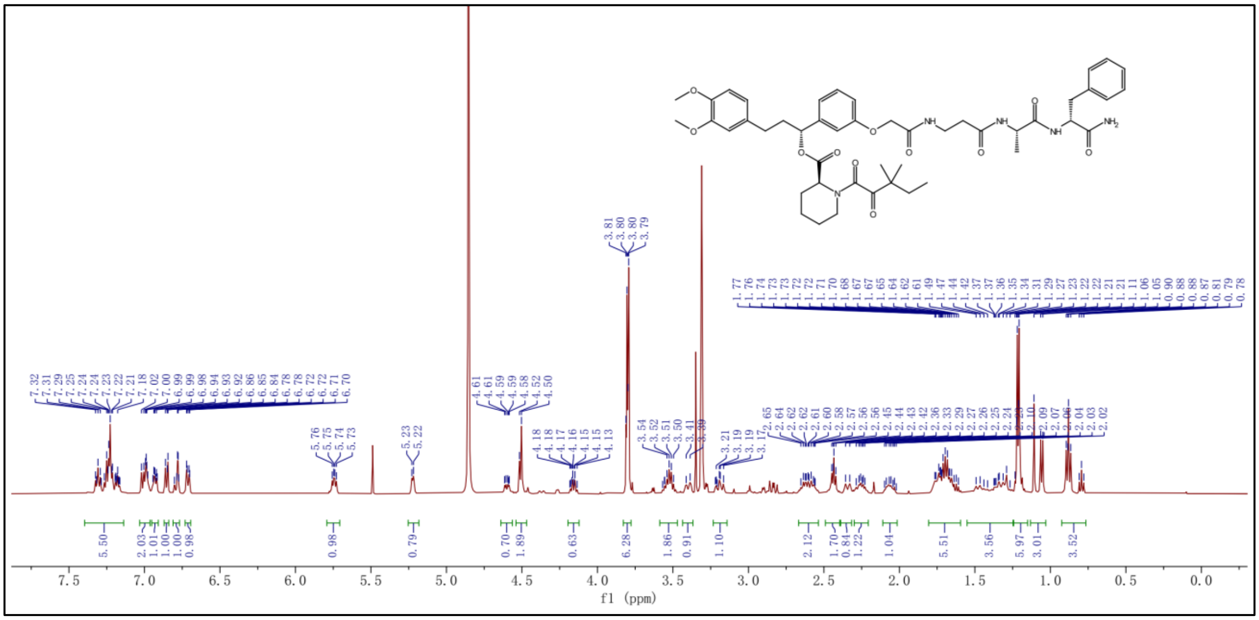

### $^{13}$C NMR spectrum of SLF-F9

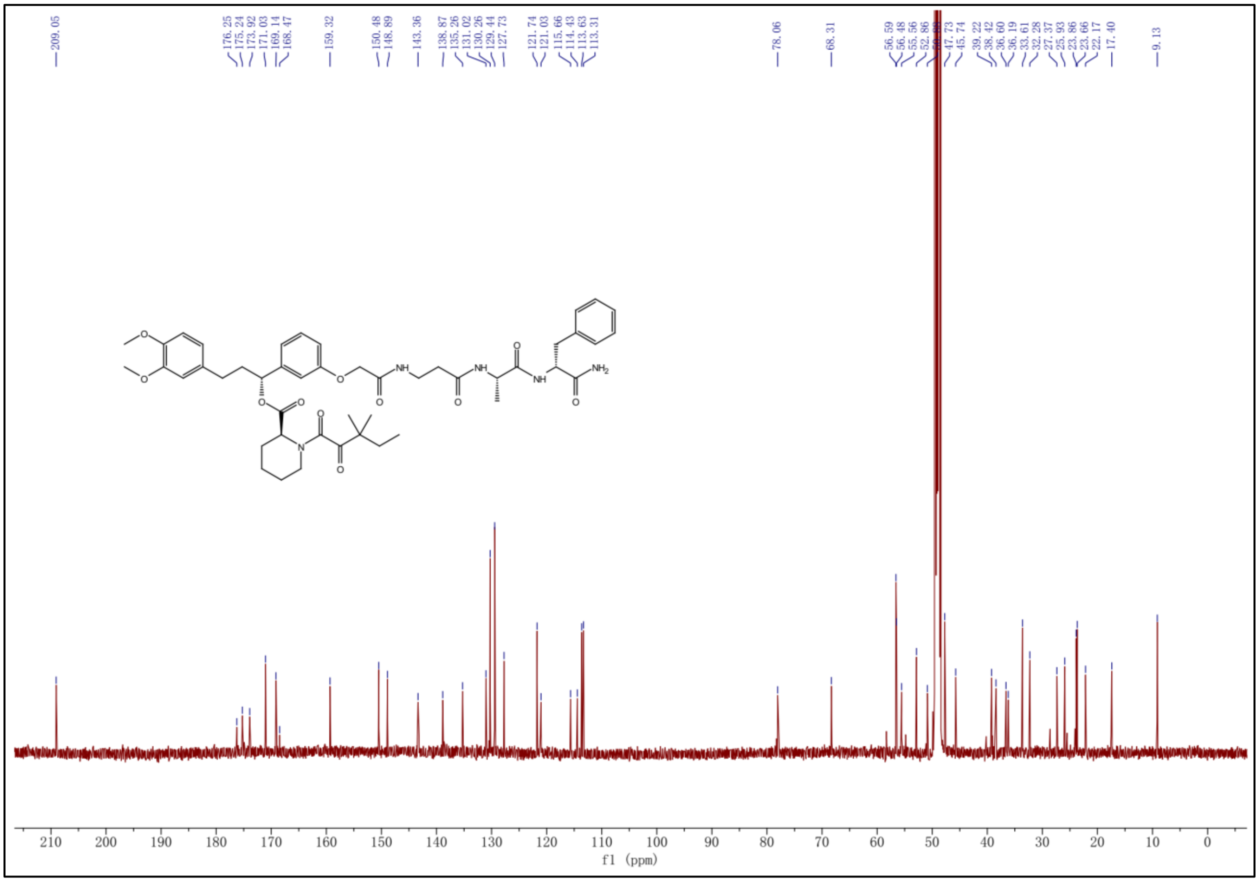

[1]H NMR spectrum of SLF-F10

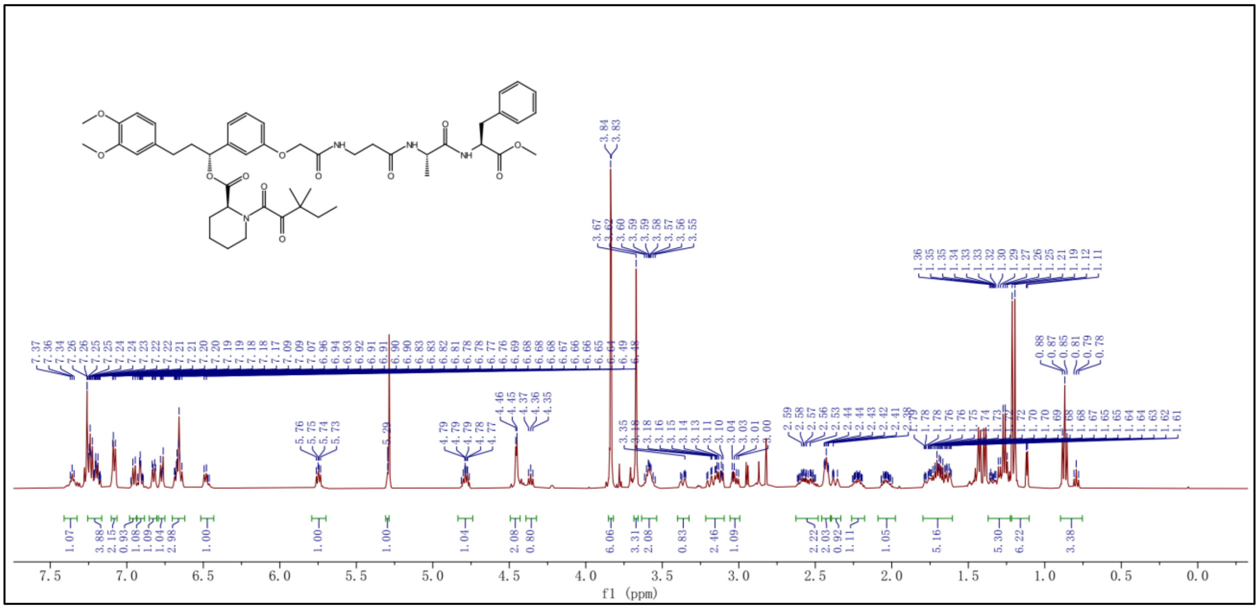

[13]C NMR spectrum of SLF-F10

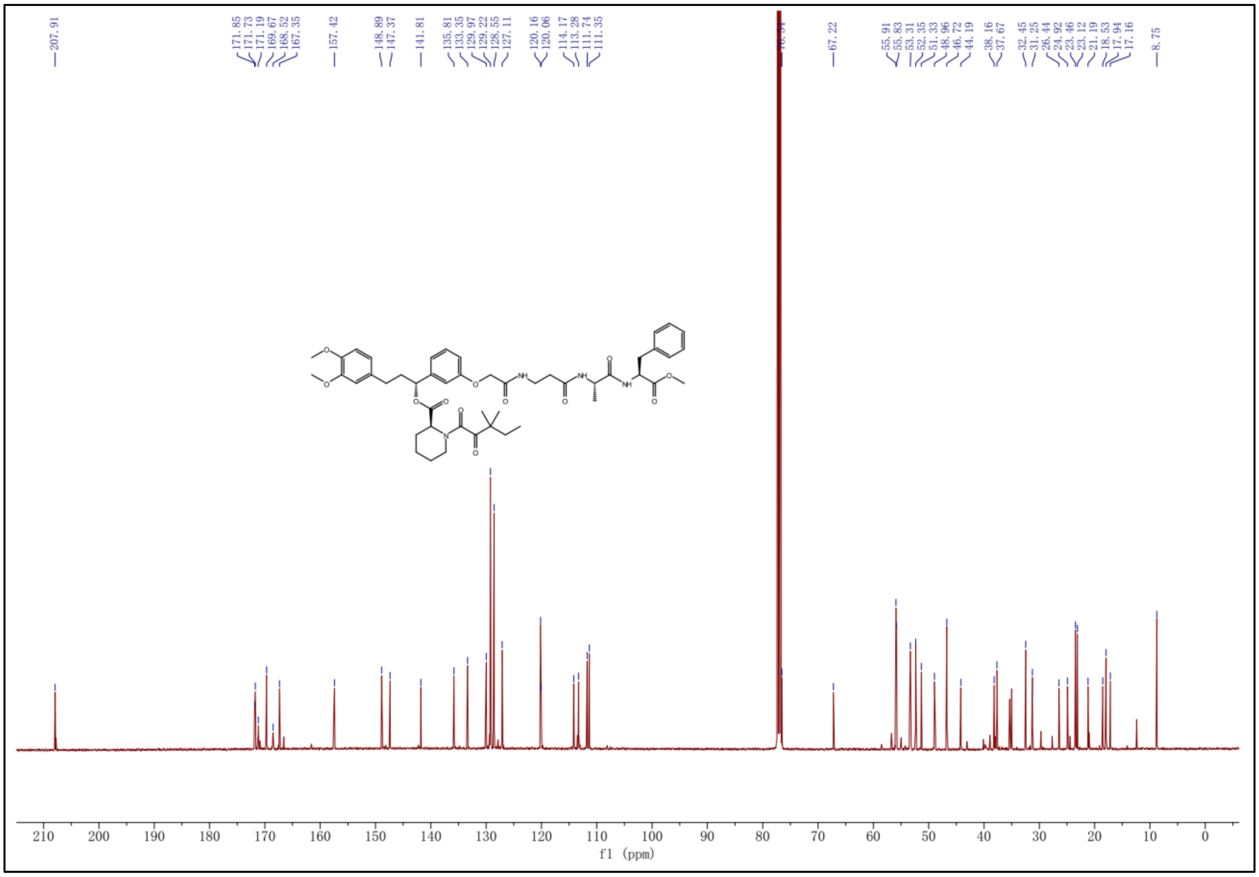

**[1]H NMR spectrum of SLF-F11**

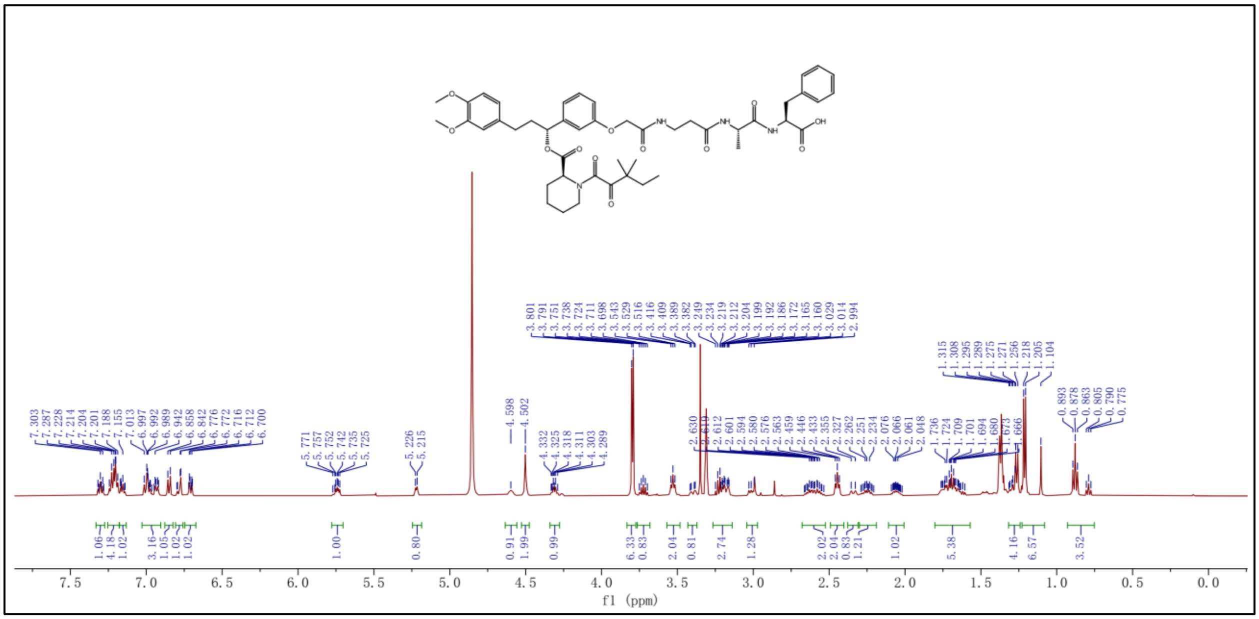

**[13]C NMR spectrum of SLF-F11**

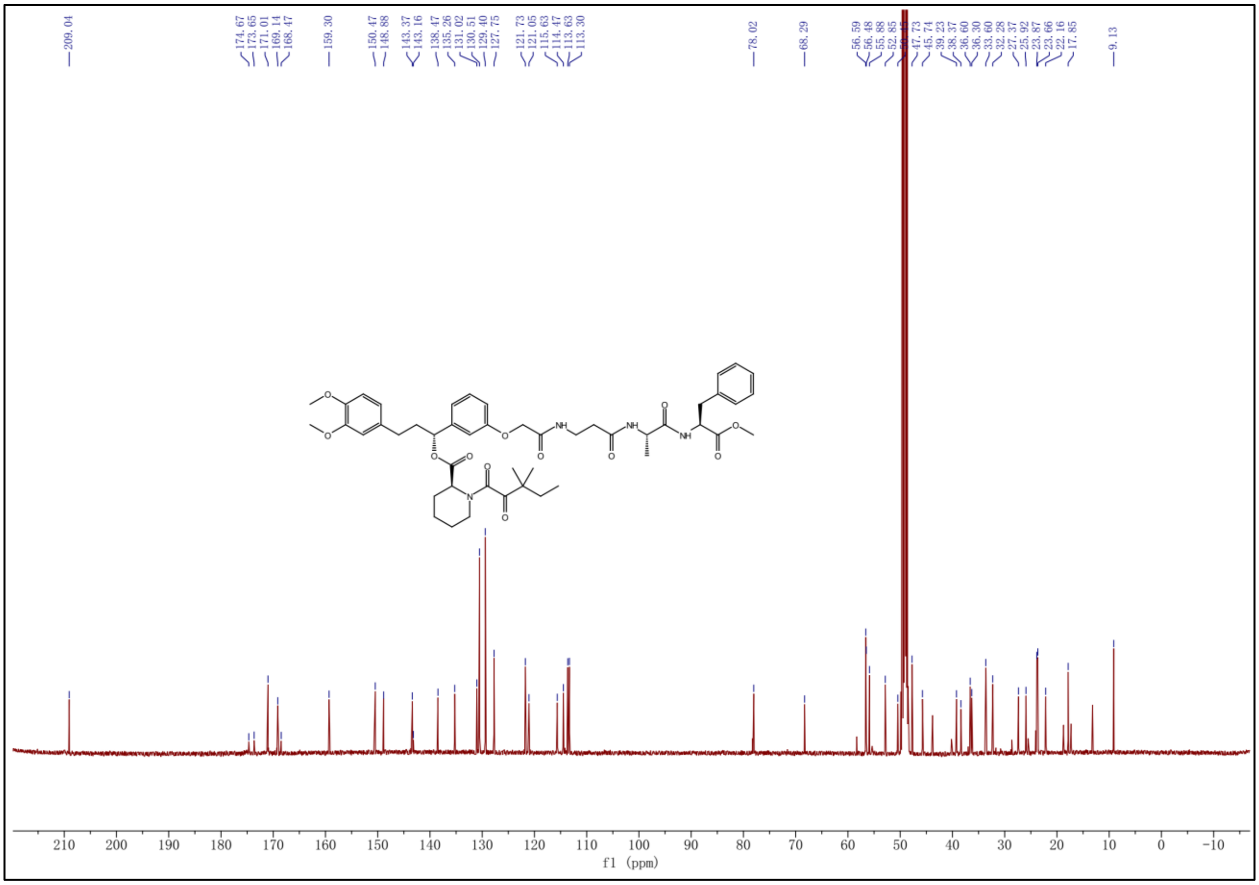

### $^{1}$H NMR spectrum of MKI-F3

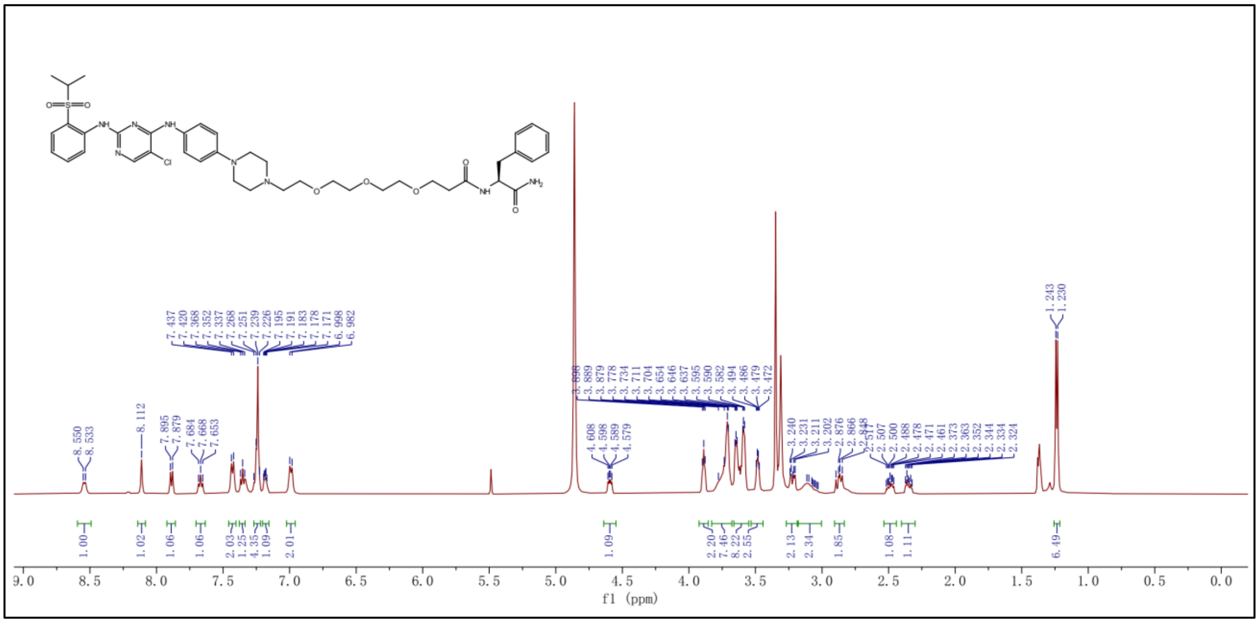


### $^{13}$C NMR spectrum of MKI-F3

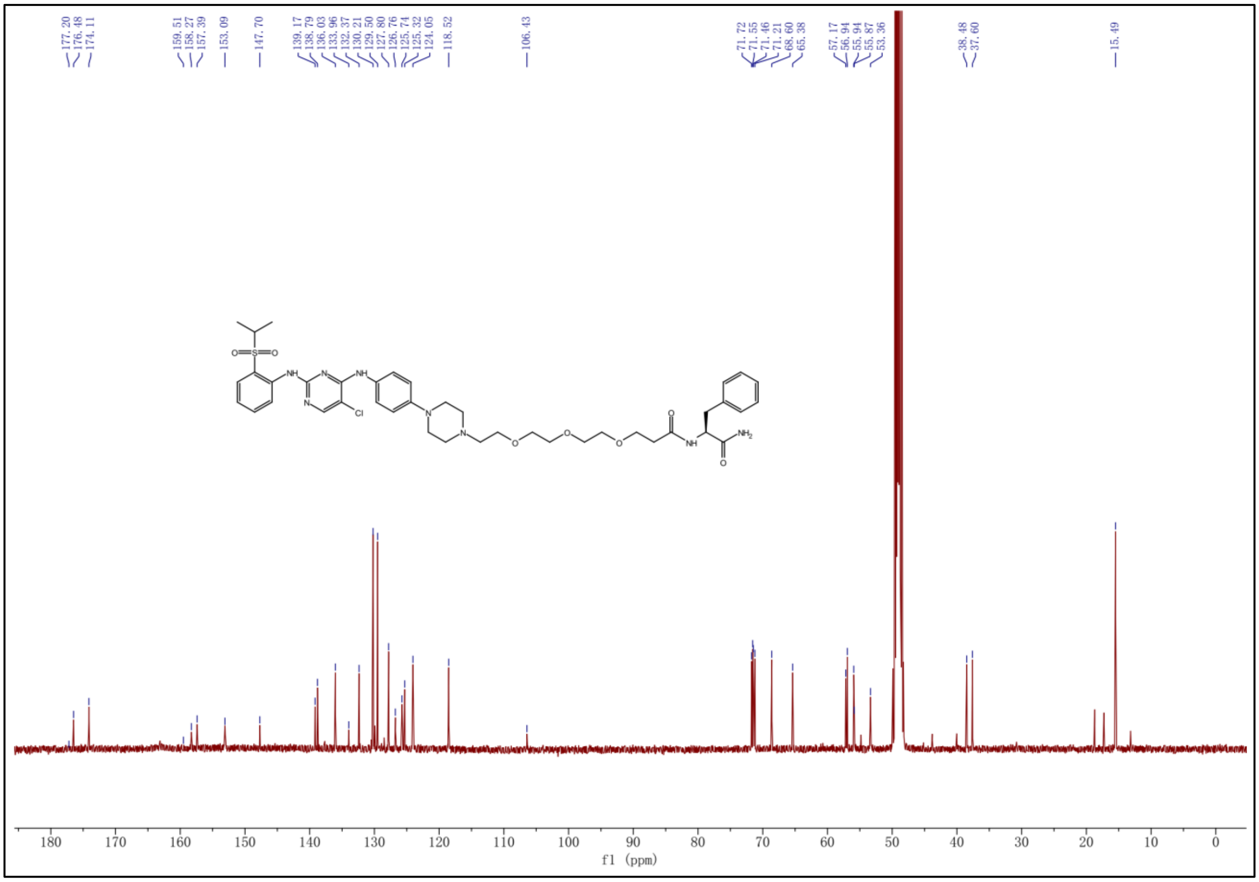

#### $^{1}H$ NMR spectrum of MKI-F4

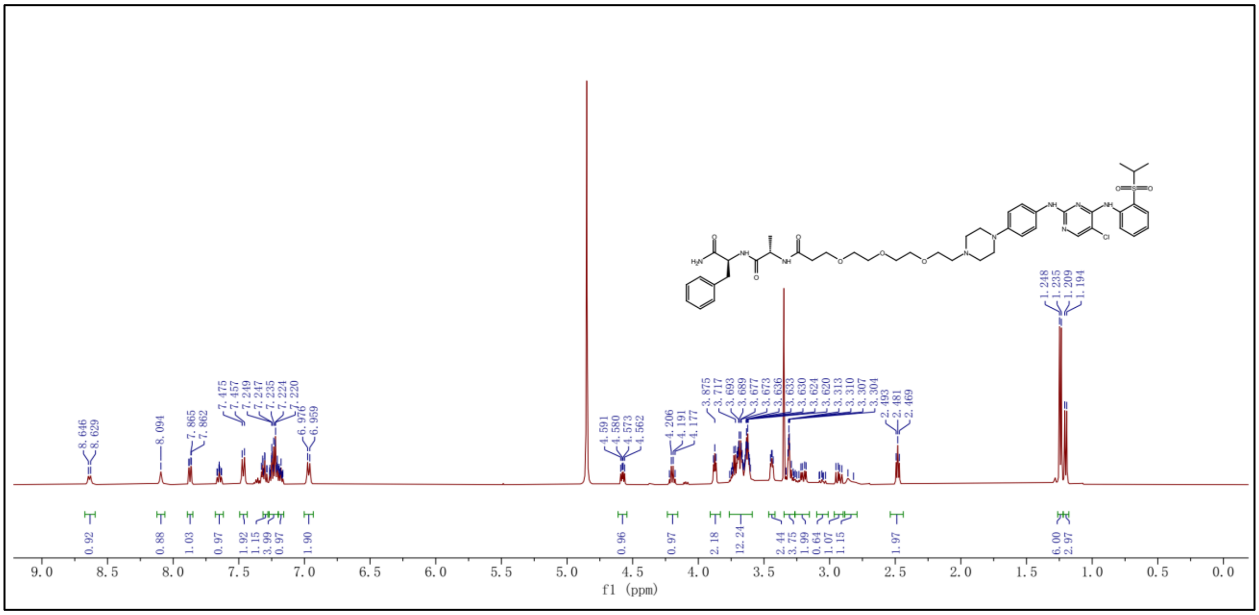

#### $^{13}C$ NMR spectrum of MKI-F4

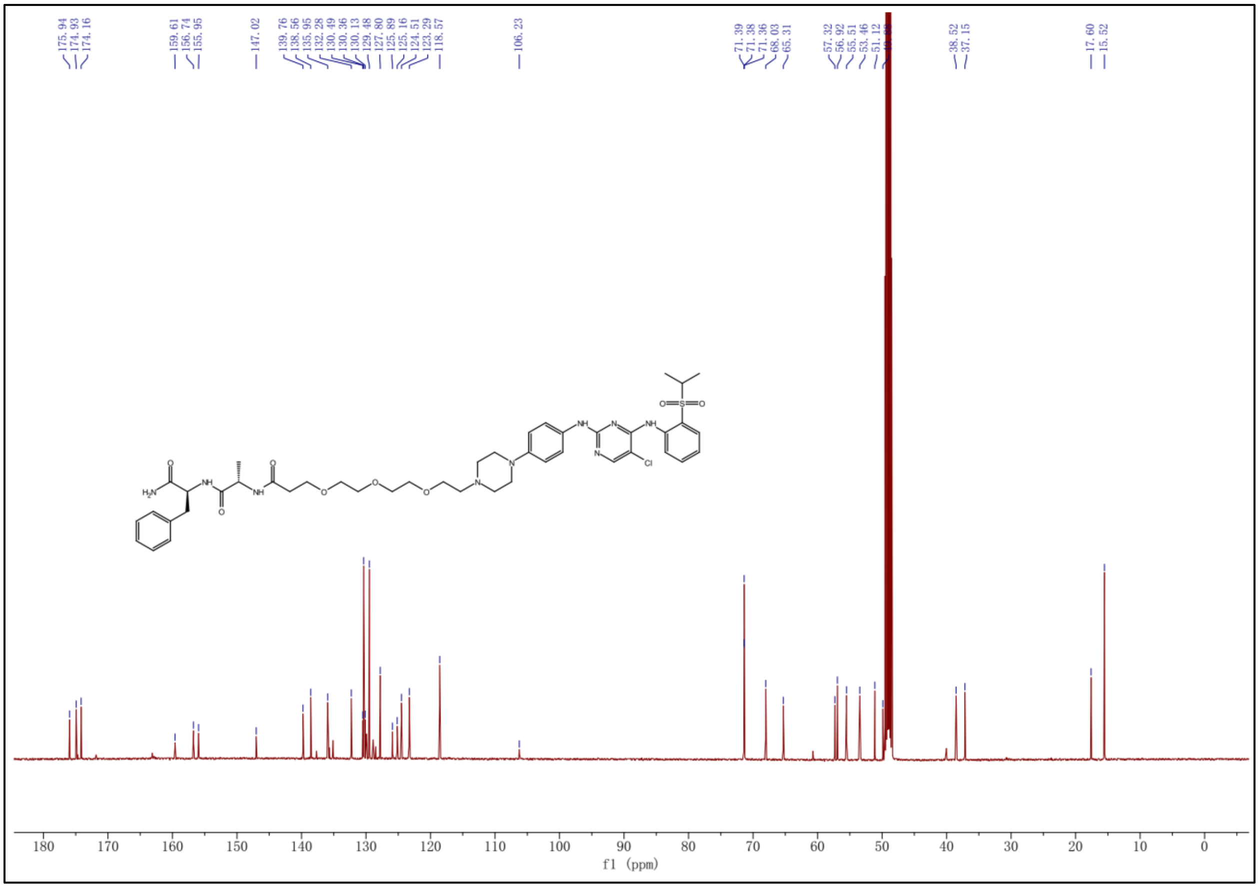

### $^{1}$H NMR spectrum of JQ1-F4

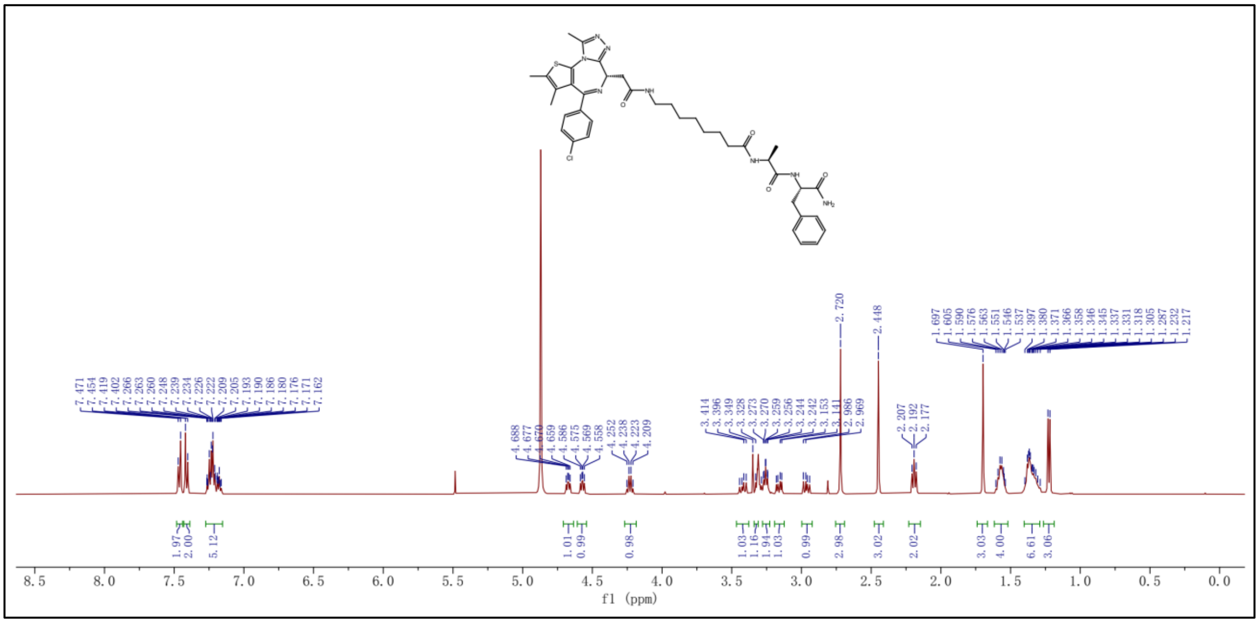

### $^{13}$C NMR spectrum of JQ1-F4

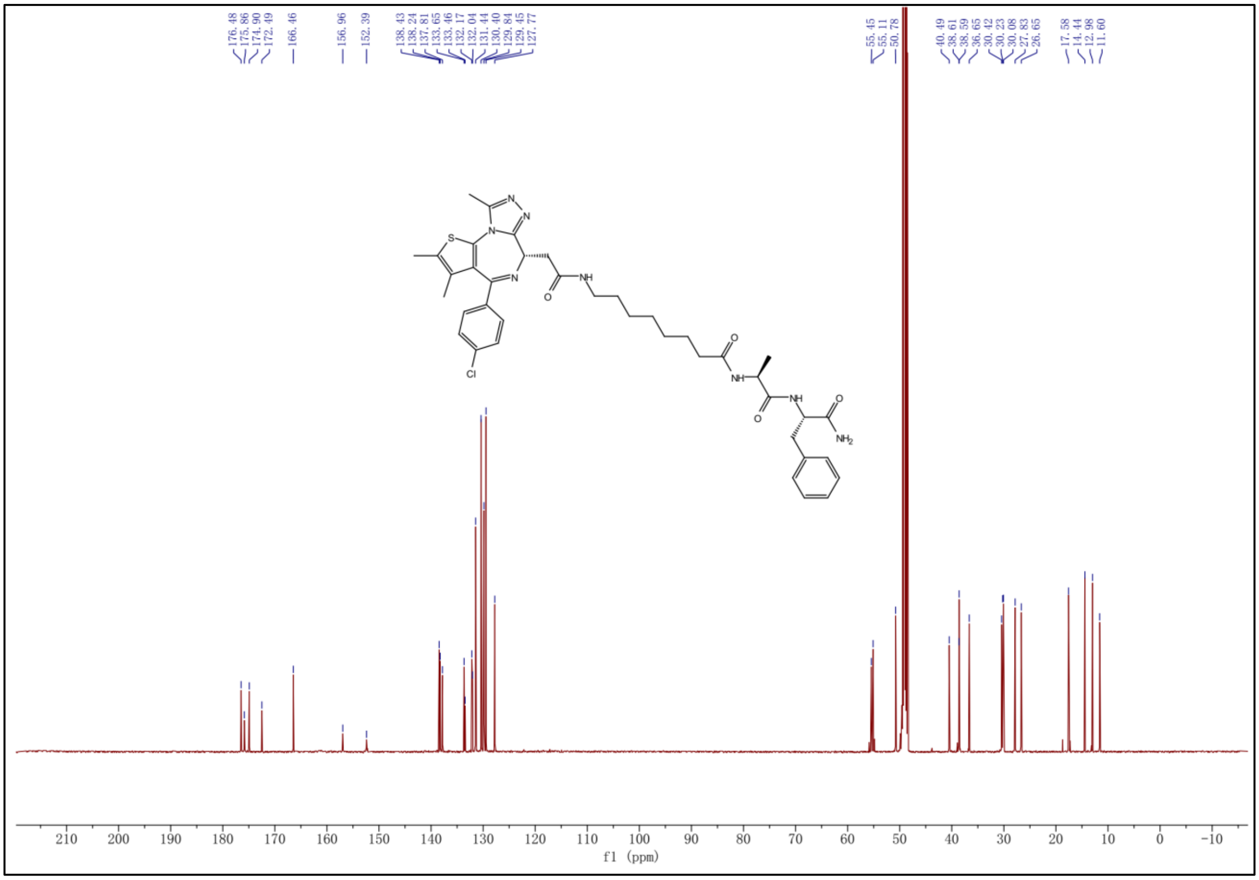
